# Evolutionary Dynamics, Evolutionary Forces, and Robustness: A Nonequilibrium Statistical Mechanics Perspective

**DOI:** 10.1101/2021.10.10.463854

**Authors:** Riccardo Rao, Stanislas Leibler

## Abstract

Any realistic evolutionary theory has to consider: *(i)* the dynamics of organisms that reproduce and possess heritable traits; *(ii)* the appearance of stochastic variations in these traits; and *(iii)* the selection of those organisms that better survive and reproduce. These elements shape the “evolutionary forces” that characterize the evolutionary dynamics. Here, we introduce a general model of reproduction–variation–selection dynamics. By treating these dynamics as a non-equilibrium thermodynamic process, we make precise the notion of the forces that characterize evolution. One of these forces, in particular, can be associated with the robustness of reproduction to variations. The emergence of this trait in our model—without any explicit selection for it—is an example of a general phenomenon, which can be called enaptation, distinct from the well-known and studied phenomena of adaptation and exaptation. Some of the detailed predictions of our model can be tested by quantitative laboratory experiments, similar to those performed in the past on evolving populations of proteins or viruses.

## INTRODUCTION

A conventional view of evolutionary dynamics is based on three essential elements [1]: *(i)* organism reproduction with imperfect heredity; *(ii)* variations, including mutations, which are typically introduced by the reproduction process; *(iii)* selection, which acts within a population and allows some variant species to survive and reproduce while eliminating others. When considering variations, a sizeable fraction of evolutionary biology is focused on genetic and epigenetic variations. However, variations upon which selection acts are occurring on multiple levels, and involve many entities, traits and behaviors that are usually encapsulated by a rather imprecise concept of phenotype [2, 3]. Regarding selection, many phenotypic aspects contribute to long-term survival and reproduction. Two instances are the interactions between the organisms (*e*.*g*., sexual reproduction, predation, competition and cooperation, social organization), and the interactions with biotic and abiotic environmental factors (*e*.*g*. the presence of other species or the inorganic composition of a certain habitat), whose changes span multiple spatiotemporal scales.

The full complexity of evolutionary dynamics is overwhelming. So, it is of no surprise that simplifying narratives have been developed to summarize its processes. The most popular one is known under the name of *adaptationism* [4]. At the basis of this narrative lies the word *adaptation*, whose etymology (from Latin *adaptāre*: *ad-*, towards + *aptāre*, to fit) evokes the emergence—through evolution—of fit organisms, *i*.*e*. organisms that can survive and reproduce in the surrounding environment. This narrative would simply be tautological, if it were not for a further elaboration on the nature of variations and their selection. Briefly stated, selection in living organisms is believed to arise from stochastic events, superimposed on “evolutionary forces”, which are the result of interactions between the organisms themselves and of their interactions with the environment. In contrast to the neutral theory of evolution [5]—which assumes that purely stochastic events dominate evolution—adaptationism states that the generation of fit organisms is mostly due to evolutionary forces.

As with any other simplifying narrative of evolutionary dynamics, pure adaptationism has found its critics. The major concern relates to its naïve premise that all observed phenotypic features have been selected for their present “aptive” (fitness) value [6]. Indeed, some of the observed phenotypic features seem to be co-opted from previously evolved traits. An often-quoted example of such functional shifts is the case of feathers: initially, they might have evolved for temperature regulation, but later were used for facilitating the flights (and water diving) of birds [7]. Such cases of preexisting features that were co-opted for new functions are now described by the term *exaptation* (from Latin *ex-*, out of + *aptāre*, to fit) [8], rather than adaptation. Obviously, the frontier between adaptations and exaptations is not sharp and thus it is being constantly disputed.

The main role of (necessarily) simple mathematical models of evolution is to analyze the possible outcomes of evolution and to explore the assumptions that generate these outcomes. In this way, one hopes to clarify some essential concepts used by evolutionary narratives. The goal of the present work is to formulate a simple model that incorporates the three essential elements described above. Consequently, we call the evolutionary dynamics described by this model the *reproduction– variation–selection* (RVS) dynamics. In formulating the model, we seek both simplicity and generality. For sake of generality, we specify neither the particular nature of the hereditary variables nor that of the related variations: they can be genetic, epigenetic, or phenotypic. For sake of simplicity, we assume large population sizes and the presence of a constant environment. Such an approach allows us to build a theoretical framework inspired by non-equilibrium statistical mechanics [9–12]. Within this framework, we can clearly define the notion of evolutionary force and explicate their relation to stochastic events. Because of the simplicity of the model, we are able to get analytical expressions for different force terms, and to perform explicit analyses of the role played by them. In particular, we uncover an evolutionary force within the RVS dynamics that can engender robustness of reproduction to variations, without any explicit selection for this trait. By adding restraining assumptions to the model, we can also make simple predictions about the behavior of population robustness during the RVS dynamics. Lastly, we compare our predictions with the results of laboratory experiments on populations of evolving viruses.

## MODEL

### Generic Evolutionary Dynamics

Generic evolutionary dynamics can be described as follows. We imagine a population of *N* organisms, each of which is characterized by some hereditary variables *γ*. These variables can describe genetic and epigenetic factors, collective phenotypic features, etc. We do not specify the precise biological nature of *γ*, but we assume that they determine the reproductive success of each organism. Mathematically, *γ* can be represented as multidimensional continuous or discrete variables, or a mixture of the two; here for simplicity’s sake we focus on the discrete case (but see [SI §II] for the continuous case). We refer to an organism characterized by the variables *γ* as a being of *type γ*.

The typical number of offspring that each type engenders in one generation is the *reproduction rate f*_*γ*_. When the environment is assumed constant, selection favors organisms reproducing faster. During one generation, random changes of the hereditary variables create the diversity upon which natural selection can act. We generically refer to these changes as *variations*. For the particular case of genetic variations, they may involve a single organism, or pairs of organisms, through recombination. As stated in Introduction, variations can also be epigenetic and/or phenotypic.

Over many generations, the evolutionary dynamics can be described as a discrete-time stochastic Markov jump process. The *population composition* is denoted by the vector *n* = (*n*_*γ*_): each of its entries represents the number of *γ* -type present in the population. The probability of observing a certain *n* at generation *τ, p*_*n*_ (*τ*), is described by a Chapman–Kolmogorov equation (see Box 1 for a descriptive summary of the main formulae),

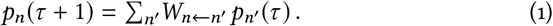

The entries of the *transition kernel W*_*n*←*n′*_ determine the probability that the population transitions from *n*′ to *n* over one generation. Notice that we assume non-overlapping generations. The precise form of *W*_*n*←*n*′_ depends on the details of the stochastic evolutionary process. In general, *f*_*γ*_, and thus *W*_*n*←*n′*_, changes with the environment.

Before committing ourselves to a particular *W*_*n* ←*n′*_, let us introduce a quantity that we call *evolutionary directionality*. It is defined as the log ratio of forward and backward transition probability,

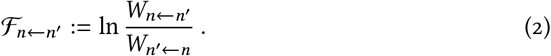

Evolutionary directionality will play a crucial role in our discussion. For a pair of transitions *n* ⇌ *n*′ such that *W*_*n*←*n′*_ ≃ *W*_*n*′ ←*n*_, neither direction is probabilistically favored. This situation corresponds to negligible directionality, ℱ_*n* ← *n′*_ ≃ 0. In contrast, when *n* ⇌ *n′* are such that *W*_*n*←*n′*_ ≫*W*_*n′*← *n*_ (respectively *W*_*n*←*n*′_*≪W*_*n′*← *n*_), the population is more likely to evolve in the direction *n*←*n*′ (respectively *n*′ ← *n*). This situation corresponds to nonzero directionality, ℱ_*n*←*n*′_ > 0 (respectively ℱ_*n*←*n′*_ < 0).

In general, the evolutionary directionality ℱ_*n*←*n′*_ can be written as a sum of two contributions, the first of which can be expressed as a *potential* difference, while the second cannot:

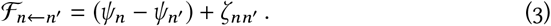

The precise form of *ψ*_*n*_ and *ζ*_*nn′*_ depends on how selection and variations affect the population, expressed in the model through *W*_*n*←*n′*_. We refer to *ζ*_*nn′*_ and *ψ*_*n*_ − *ψ*_*n′*_ as *evolutionary force contributions*, and they are important for the following reason. If *ζ*_*nn′*_ is positive for *n* ← *n′*, then *ζ*_*nn′*_ positively contributes to the directionality ℱ_*n*←*n′*_ and we can say that this force contribution favors such transition. Analogously, if *ψ*_*n*_ increases along *n*←*n*′, then *ψ*_*n*_ − *ψ*_*n′*_ is positive and such a transition is favored by the force contribution originating from *ψ*_*n*_.

In analogy to similar quantities encountered in nonequilibrium statistical mechanics and thermodynamics, see *e*.*g*. [13], we refer to the two contributions in Eq. (3) respectively as *conservative* and *nonconservative* evolutionary forces. Note that when the latter force vanishes, *ζ*_*nn′*_ = 0 for all *n* and *n′*, one can show that 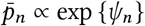 is the stationary distribution of the dynamics described by (1), see *e*.*g*. [14]. This situation corresponds to a gradient stochastic dynamics in which the landscape identified by *ψ*_*n*_ determines the conservative (or gradient) force *ψ*_*n*_ − *ψ*_*n′*_.

### Reproduction–Variation–Selection Dynamics

A model of evolutionary dynamics is specified by a particular choice of the kernel *W*_*n*←*n′*_. In our RVS dynamics, this kernel is defined through the probability that a variation *γ* ← *γ*′ occurs over a single generation, *π*_*γ*←*γ*′_ (where *π*_*γ*←*γ*′_ ≥ 0 and ∑_*γ*_ *π*_*γ*←*γ*′_ = 1 for all *γ*′). We consider reversible variations, so that *π*_*γ*←*γ*′_ > 0 ⇔ *π*_*γ*′ ← *γ*_ > 0. This assumption does not preclude that variations in some of the directions may be much more likely than in the opposite ones, *i*.*e*. *π*_*γ*←*γ*′_ ≫ *π*_*γ*′ ← *γ*_ for some *γ* and *γ*′ Crucially, *π*_*γ*←*γ*′_ must be regarded as a function of *γ*′. Indeed, *π*_*γ*←*γ*′_ is also subject to evolution: different types may evolve to vary in different ways. In general, many detailed mechanisms can contribute to variations of a type, all encapsulated in the probability *π*_*γ*←*γ*′_. In addition, for variations involving pairs of organisms, the corresponding contribution to *π*_*γ*←*γ*′_ depends on the population composition *n*. For the sake of simplicity, here we do not distinguish among different mechanisms of variation (see [SI § IE] for such details). We consider only the overall probability that *γ* has varied:

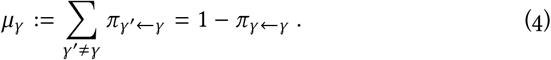

In our model, the creation of a new pool of variants and the selection from this pool take place in each generation, as in Fig. 1. We assume the overall size of the population to be large, *N* = ∑ _*γ*_ *n*_*γ*_ ≫ 1, so that its fluctuations can be neglected and its value regarded as a constant [SI §IB]. This might reflect the fact that environmental resources are abundant but limiting. For simplicity’s sake we also assume that the organisms do not directly interact, so that the reproduction rate *f*_*γ*_ does not depend on the population composition *n*. The case of interacting populations with small and fluctuating population sizes is analyzed in [SI §IA].

**FIG. 1.**
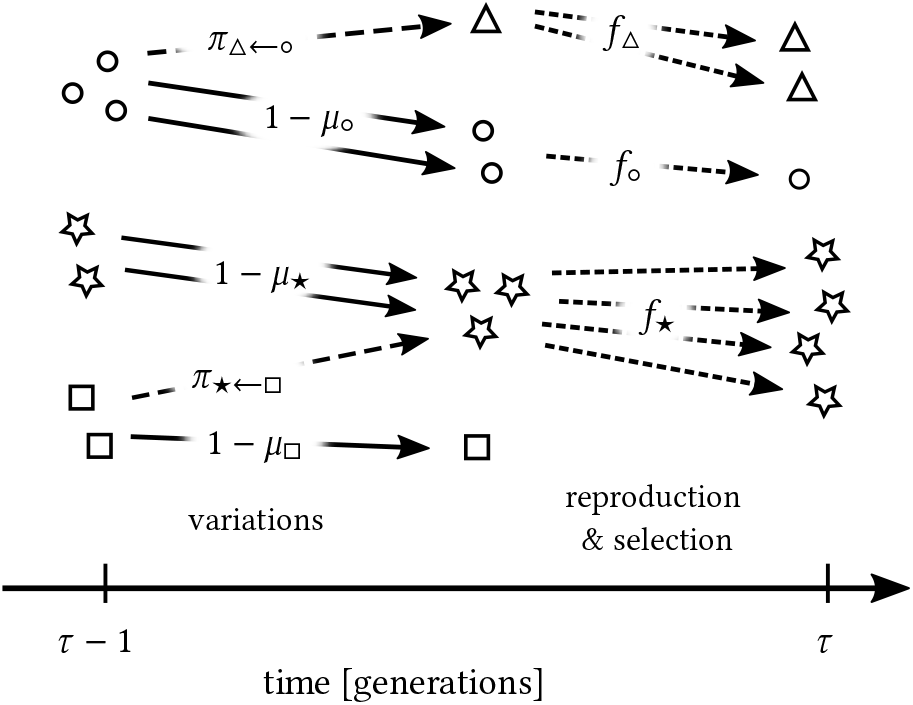
Schematic representation of the evolutionary dynamics as modeled by the transition probabilities in Eq. (5). The symbols ○, ⋆, and □ represent different types *γ* present at generation *τ* − 1, in a population of *N* = 7 organisms. As variants are generated—dashed lines—a new type appears, Δ. Continuous lines denote no variation. The variation probabilities *π*_*γ* →*γ′*_, as well as the probability that no variation occurs, 1 − *μ*_*γ*_, are also reported. Selection—dotted lines— finally determines which types and in what amount make it to the next generation. The reproduction rate *f*_*γ*_ determines the chances of being selected.

Under these assumptions the transition kernel can be expressed in a simple form (details in [SI §§IA–C]),

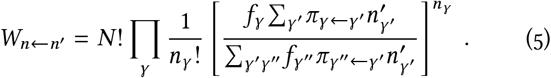

Note the alternation of variations, reproduction and stochastic selection, as in Fig. 1. The term in square brackets describes *(i)* the probability that variations affect the organisms, 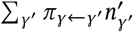, and *(ii)*organismal reproduction, *f*_*γ*_. Then, the multinomial product 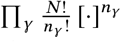 ensures the random selection of *N* organisms. Fast growing types—types with larger numbers of descendants, *f*_*γ*_ —are more likely to be selected, yet *selection fluctuations* (usually referred to as *genetic drift* in the literature of evolutionary dynamics) are accounted for. The model (5) can be regarded as a generalized version of the Wright–Fisher model [15] in which types and variations are generic, and types are subject to selection.

## RESULTS

### Evolutionary Forces

For the model described by the kernel (5), the conservative force potential and the nonconservative force contribution are

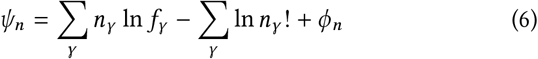

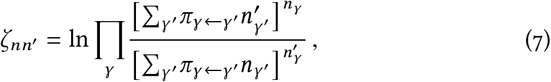

where

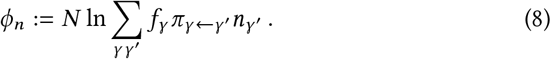

As discussed in [SI §IC], these expressions follow from a simple algebraic procedure using (5). This general procedure is justified by the fact that we do not specify the nature of *γ*, nor how variations affect *γ*.

We can now analyze the expressions for *ψ*_*n*_ and *ζ*_*nn′*_ and find conditions for which *ζ*_*nn*′_ is positive, and for which *ψ*_*n*_ increases. As mentioned before, such conditions indicate compositions *n* towards which the population is more likely to evolve.

The nonconservative force *ζ*_*nn*′_ accounts for the effect of variations and selection fluctuations (a.k.a. genetic drift). This fact is highlighted by the presence of the variation probabilities *π*_*n*←*n′*_ and the multinomial product in Eq. (7). Although in general one cannot determine easily the sign of *ζ*_*nn′*_, two limiting cases are useful to gain some insight into the effect of this nonconservative force (details in [SI §§III and ID]). We first consider the limit of small variation probabilities (4), *μ*_*γ*_ ≃ *μ* → 0. Imagine a population composition transition *n*←*n*′ in which a particular type *γ* disappears: 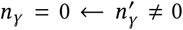. This event is irreversible since the nonconservative force diverges logarithmically (*ζ*_*nn′*_ = − *ζ*_*n′n*_ ∼ − ln *μ* → ∞, for *μ* → 0), thus preventing *γ* reappearance. Generalizing this insight to small but finite *μ*_*γ*_, we can argue that *ζ* typically causes a loss of diversity by favoring the selection of the most abundant types. Second, consider the idealized case when the variation probability solely depends on the variant type, *i*.*e. π*_*γ ← γ*′_ = *π*_*γ*_ for all *γ* and *γ*′. The probability *π*_*γ*_ quantifies the likelihood that *γ* is engendered by variations, irrespective of the type from which it originates. In this case, the nonconservative force actually becomes conservative, *ζ*_*nn′*_ = ∑_*γ*_*n*_*γ*_ ln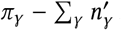 ln *π*_*γ*_. It generically favors *n* with higher ∑_*γ*_ *n*_*γ*_ ln *π*_*γ*_, *i*.*e*. it favors population compositions with higher cumulative log likelihood of being engendered.

The potential *ψ*_*n*_ that appears in the conservative force is composed of three terms (Eq. (6)). The first term is associated with selection, since it is simply the cumulative logarithmic growth of the population. The corresponding evolutionary force, 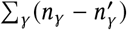 ln *f*_*γ*_, drives the population towards compositions with higher cumulative log reproduction rate.

The second term is an entropic contribution, and it arises due to the assumed indistinguishability of organisms of the same type. The corresponding force, ∑_*γ*_ ln 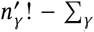 ln *n*_*γ*_!, favors heterogeneous population compositions. As a limiting case, consider again a transition in which a particular type *γ* disappears: 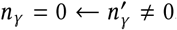. Such an event is disfavored by this entropic force, which scales as the logarithm of the population size (∼ − ln *N* < 0). Hence, for constant *μ* and *μN* > 1, the entropic force counteracts the loss of diversity generated by the nonconservative force *ζ* (details in [SI §III, Eq. (S51)]).

Finally, the last term in Eq. (6), Eq. (8), can be associated with the combined effect of selection and variations. We name this term *expected growth potential* of the population *n*, since it quantifies the expected growth of *n* at the next generation. It is larger for populations that are likely to generate variants with high reproduction rate. This kind of population is thus favored by the force term

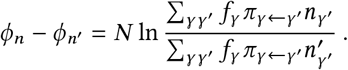

Since *ϕ*_*n*_ depends on *n* in a nonlinear fashion (see the logarithm), *ϕ*_*n*_ can be viewed as describing an effective interaction acting among the types present in the population. This interaction is engendered by the combined action of selection and variations.

All main mathematical expressions are summarized in Box 1, whereas the nature and effect of each evolutionary dynamical force ((6) and (7)) are illustrated in Box 2 using two simple toy models.

### Robustness

Robustness is an important quantity that measures the extent of insensitivity of an organism, or its particular phenotypic features, to extrinsic or intrinsic variations. For instance, the so-called mutational robustness measures how much a given phenotypic feature changes upon genetic mutations (in a given environment). The expected growth potential *ϕ*_*n*_ can be explicitly related to robustness. To see this, let us first introduce *ω*_*γ*_, a measure of the *sensitivity* (i.e. the inverse of robustness) of the reproduction rate to type variations,

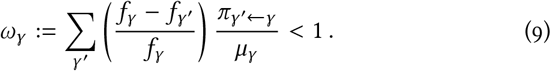

Types *γ* with sufficiently large reproduction rate *f*_*γ*_ have positive sensitivity, *ω*_*γ*_ > 0, see also Fig. 2. Large values of *ω*_*γ*_ (*ω*_*γ*_ ≲ 1) imply that the relative decrease of the reproduction rate, (*f*_*γ*_ − *f*_*γ*′_)/ *f*_*γ*_, averaged over all possible variations, *π*_*γ*′ *← γ*_*/μ* _*γ*_, is large. Hence, types with large *ω*_*γ*_ have large sensitivity, or small robustness to variations. In contrast, *ω*_*γ*_ ≃ 0 characterizes those types whose reproduction is insensitive— and hence robust—to variations.

**FIG. 2.**
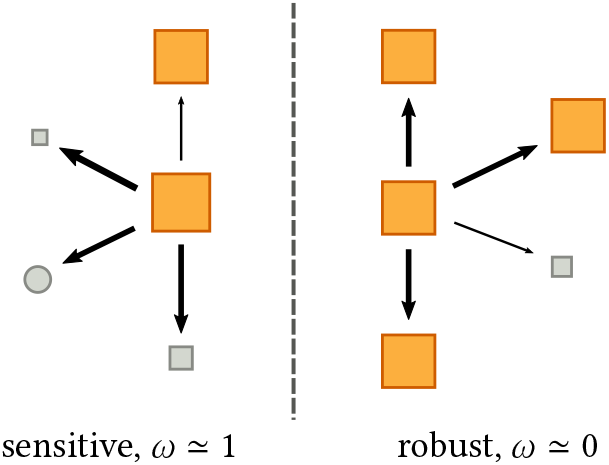
Pictorial illustration of sensitivity. Each square represents a type: fast-reproducing types are represented by large orange squares, whereas slow-reproducing ones by small gray squares. Arrows identify possible variations, and their thickness reflects the probability of variations. Consider two types with high reproduction rate, central orange squares. (Left) A sensitive type is very likely to vary into a type with lower reproduction rate. (Right) A robust type vary with high probability into types with similarly high reproduction rate.

The expected growth potential *ϕ*_*n*_ can be now expressed as a decreasing function of *ω*_*γ*_ :

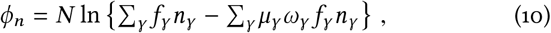

*i*.*e*. the lower the sensitivity of the organisms of the population, the higher *ϕ*_*n*_, see [SI §IE].

To gain further understanding of Eq. (10), let us consider populations 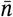 composed of types with relatively high and roughly equal reproduction rates: *f*_*γ*_ ≃ *f*_high_, for any *γ* such that 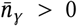. This case could describe situations in which selection is so strong that only the fastest-reproducing types survive. Equation (10) can be approximated as

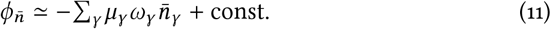

(see [SI §IE]). The right hand side can be interpreted as the cumulative variation probability times sensitivity of the population 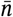. Hence, there are only two ways in which the potential term 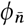 can increase: either the types of the population can be subject to less variations, *i*.*e. μ*_*γ*_ decreases, or they can be more robust to variations, *i*.*e*. their sensitivity *ω*_*γ*_ decreases.

Equations (10) and (11) have thus a general, important significance: they show how the evolutionary force generated by *ϕ*_*n*_ − *ϕ*_*n′*_ favors populations whose types exhibit either low variation rates, *μ*_*γ*_ ≪ 1 or high robustness to variations, *ω*_*γ*_≪ 1. Crucially, the emergence of these features is a property of generic reproduction–variation–selection dynamics and is not restricted to either specific hereditary variables or specific variation mechanisms.

#### Box 1

Summary of Mathematical Expressions

**RVS Dynamics**

Chapman-Kolmogorov for a generic evolutionary dynamics

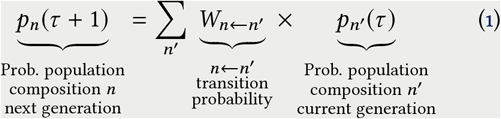

Evolutionary directionality

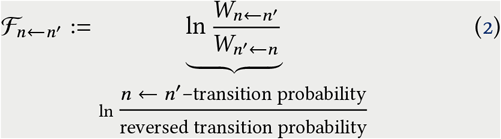

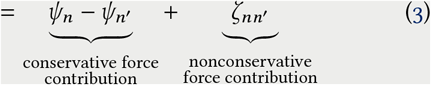

RVS Dynamics – Evolutionary Forces: Nonconservative force

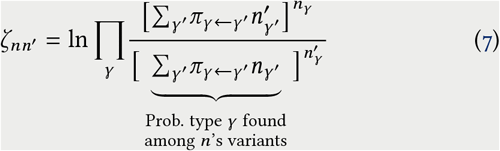

RVS Dynamics – Evolutionary Forces: Potential

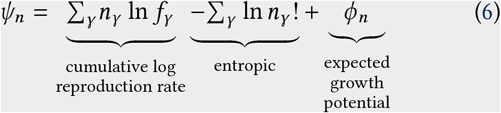

RVS Dynamics – Expected Growth Potential

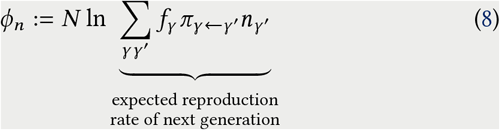

Important limiting case: 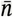 such that

- *f*_*γ*_ ≃ *f*_high,_ for any *γ* such that 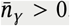.

Expected Growth Potential

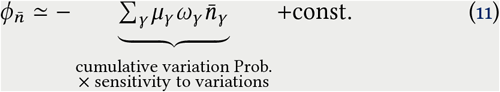

where *μ*_*γ*_ and *ω*_*γ*_ are given in Eq.(4) and (9).

**Restricted RVS Dynamics**—Additional assumptions:

- Strong selection (see SI);
- Type-independent and small variation Prob. *μ*_*γ*_≃ *μ*.

Expected Growth potential

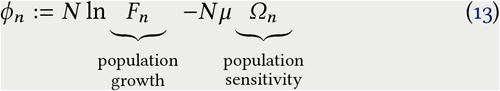

Fluctuation Relation

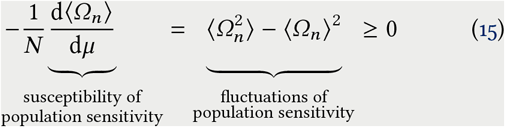

where *F*_*n*_ *:=* ∑ _*γ*_ *f*_*γ*_ *n*_*γ*_ and *Ω*_*n*_ is given in Eq.(14).

### Restricted RVS Dynamics and Fluctuation Relation

Equations (3), (6), and (7) specify what are the forces driving the evolution of RVS dynamics and how these forces shape its evolutionary outcome. However, the stochastic dynamics described by Eq. (1) and (5) cannot be solved exactly. Hence, in general, it cannot be rigorously assessed to which extent each force contributes to the evolutionary outcome. To partially overcome this limitation, we will now discuss a simple yet biologically relevant case, for which we do have an approximate analytical solution.

First, we neglect the nonconservative force *ζ*_*nn′*_ (7). Heuristically, this can be justified when the variation probabilities *μ*_*γ*_ s are not too small and selection is strong compared to selection fluctuations—types are highly discriminated by their reproduction rate, see [SI §IV]. These conditions guarantee that *ζ*_*nn′*_ remains finite and that the effect of the forces that involve selection predominate over *ζ*_*nn′*_ (we recall that *ζ*_*nn′*_ may diverge for small *μ* and does not account for reproduction and selection). When neglecting *ζ*_*nn′*_, the dynamics becomes approximately conservative, and *p*_*n*_ (*τ*) converges to a Boltzmann-like probability distribution—provided that *W*_*n* ← *n′*_ (5) is ergodic—

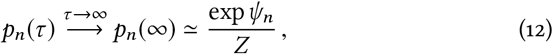

where *Z* := ∑_*n*_ exp *ψ*_*n*_ is a normalization factor. For long times, population compositions with higher *ψ*_*n*_ are exponentially more likely to be observed.

Second, we limit our analysis to a regime in which the variation probability is small and similar for all types: *μ*_*γ*_ := ∑_*γ*′ ≠ *γ*_ *π*_*γ*′←*γ*_ ≃ *μ*, for all *γ*. We can hence approximate the conservative potential (6) as (details in [SI §IV])

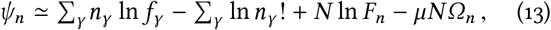

where *F*_*n*_ := ∑_*γ*_ *f*_*γ*_ *n*_*γ*_ is the cumulative growth, and

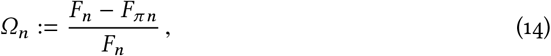

with *F*_*π n*_ := ∑_*γ, γ*′(≠*γ*)_ *f*_*γ*′_*π*_*γ*′←*γ*_ *n*_*γ*_ /*μ* being the cumulative growth upon variations. In analogy with the sensitivity of the type *ω*_*γ*_, Eq. (9), *Ω*_*n*_ can be viewed as the sensitivity of the whole population. *Ω*_*n*_ is also akin to so-called *variation loads*, where *F*_*n*_ is replaced by some maximal or reference population reproduction rate, see *e*.*g*. [16, 17].

In the asymptotic equilibrium regime (12), the population behaves similarly to a thermodynamic system in equilibrium with the environment. The population sensitivity *Ω*_*n*_ can be regarded as an energy function and *μN* as an inverse temperature. Since *ψ*_*n*_ depends in a nonlinear way on *n*, the detailed properties of the distribution (12) remain difficult to assess. However, a simple calculation inspired by equilibrium statistical mechanics (see [18] and [SI §IV]) allows us to gain insights about how *p*_*n*_(∞) affects the population sensitivity *Ω*_*n*_. Indeed,

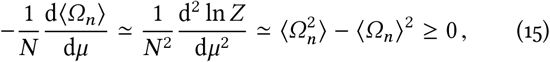

where ⟨*Ω*_*n*_⟩ = ∑_*n*_ *p*_*n*_(∞)*Ω*_*n*_ denotes the average sensitivity over the equilibrium distribution. This relation binds the fluctuations of the sensitivity (r.h.s.) to the changes of its average (l.h.s.). In the context of thermodynamics, d⟨*Ω*_*n*_⟩ /d*μ* are called *susceptibilities* as they quantify how a certain macroscopic observable, here ⟨*Ω*_*n*_⟩, is susceptible to the changes of a certain parameter, here *μ*. Equation (15) tells us that the fluctuations of *Ω*_*n*_ are higher when ⟨*Ω*_*n*_⟩ is more susceptible to changes of *μ*.

There are three important implications of the fluctuation relation (15). First, the average population sensitivity ⟨*Ω*_*n*_⟩ decreases as a function of the variation probability, d ⟨*Ω*_*n*_⟩ /d*μ* ≲ 0. Since the inverse of ⟨*Ω*_*n*_⟩ measures the population robustness to variations, this result shows that robustness increases with *μ*. Second, the fluctuations of sensitivity vanish for *N* → ∞. Third, assuming that ⟨*Ω*_*n*_⟩ saturates at high values *μ* reaching a plateau, the fluctuations 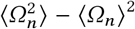 are expected to decrease with *μ*.

The fluctuation relation (15) is the main result obtained with additional assumptions imposed on the general RVS model. Although this relation has been derived with additional assumptions and is only approximate, it seems consistent with published results of laboratory experiments performed on viral populations (see Discussion and [SI §VIII]). Comparison with numerical simulations are discussed in Box 2 and in [SI §V].

## DISCUSSION AND CONCLUSIONS

From a physicist’s point of view evolution is an example of a non-equilibrium stochastic dynamical system [19–24]. One of the main reasons why it is very different from dynamics studied in physics or chemical physics is that the reproduction of organisms is tightly connected with relatively precise, longtime inheritance (in contrast to self-reproduction of simple molecular systems, such as amphiphilic micelles [25]). This allows the existence of stable variants that can better survive and reproduce than others, and thus allows organisms to evolve. Since the so-called evolutionary Modern Synthesis [26, 27] was deeply anchored in population genetics [28], mathematical models of evolutionary dynamics mostly dealt with genetic variations.

### Evolutionary Forces Clarified

In this work, we have tried to move away from this dominant scheme of modeling of evolutionary processes. Our reproduction–variations–selection dynamics model is closely related to stochastic models of nonequilibrium thermodynamic processes [9–12]. There are no assumptions about the nature of either the hereditary variables or the variations: they can be genetic, epigenetic or phenotypic. The attractive aspect of our simple model is that it clarifies the notion of “evolutionary forces”. These forces emerge from stochastic variations and selection. They can be divided into two classes: “conservative” ones, that can be expressed in terms of potentials, and “nonconservative” ones, that cannot, Eq. (3), [13, 29]. Non-conservative forces, well-studied in many physical systems, have not been explored until recently in evolutionary biology. Their effect is not intuitive, since a usual “landscape metaphor” cannot be easily applied to them [30]. This is not the case for the conservative forces, which can be easily intuited. In our model we find three separate terms which contribute to the conservative force, Eq. (6). The first term simply describes selection for fastest reproducing variants, while the second is purely entropic and induces population diversification. The remaining third conservative force contribution can be related to reproduction robustness of individual types, Eq. (10). The existence of this “robustness-generating” evolutionary force is in fact a somewhat surprising result of our analysis.

Our dynamical analysis of evolutionary forces also clarifies an academic controversy about the interpretation of these forces [31]. It was indeed debated whether selection, variations, and selection fluctuations should be interpreted as Newtonian forces (*viz*. causes of changes), or as statistical pseudo-forces (*i*.*e*. stochastic events which affect the population). From our analysis, it becomes clear that these three elements should be interpreted as pseudo-forces, but nonetheless they combine into Newtonian forces, whose expression appears in Eqs. (3), (6), and (7).

#### Box 2

Illustrative Toy Models

We consider here two examples whose purpose is to illustrate the expression of the evolutionary forces, the enaptation of robustness, and the fluctuation relation.

*Two-types Model: Nature of Evolutionary Forces* Consider an RVS dynamics with three organisms, *N* = 3, and two types, *γ*_1_ and *γ*_2_. Both types vary with the same probability, 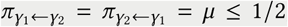, and—without loss of generality— 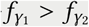. The network of type variations and that of transitions in population space are

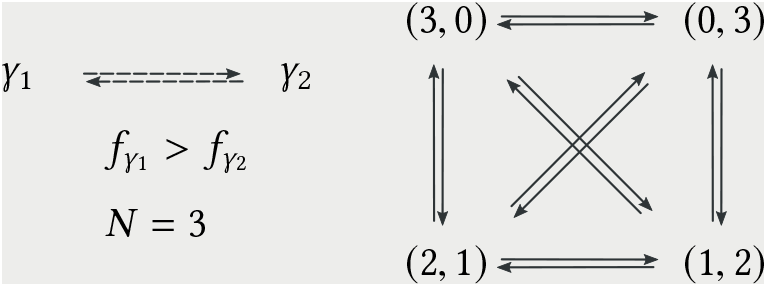

In the latter space, the reproduction rate and entropic forces act respectively as follows (details in [SI §VI])

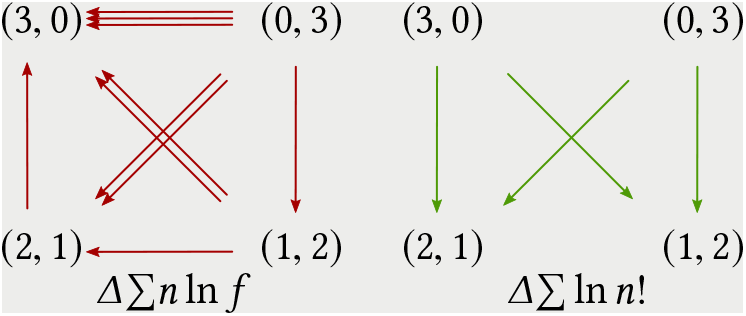

where the number of arrows is proportional to the strength of the force. The reproduction rate force favors the population with the highest proportion of fast-reproducing types, (*n*_1_, *n*_2_) = (3, 0), while the entropic force favors a diverse population composition, (*n*_1_, *n*_2)_ ∈ {(1,2), (2,1)}. Note that, since both of these forces are conservative, the sum of their values along any cycle vanishes. In contrast, the nonconservative force favors homogeneous compositions, and the sum of its values along cycles does not vanish, in general,

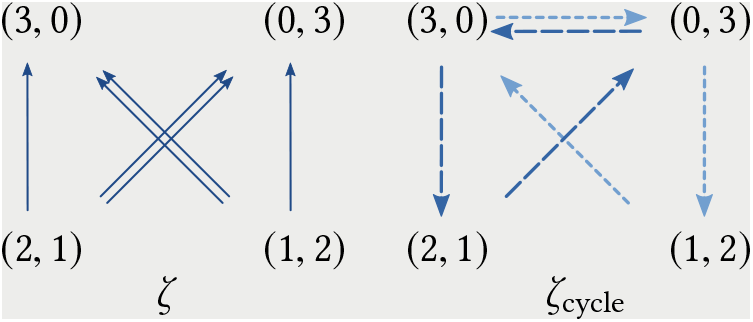

This force is stronger on the diagonal transitions (double arrows) than on the other ones. This can be explained by the fact that the loss of types (“loss of diversity”) is higher: two individuals of a certain type are lost rather than one (vertical transitions, single arrow) or none (horizontal transitions, no arrow). The right panel depicts how this force acts along cycles. See [SI §VI] for mathematical details and a discussion about the force arising from the expected growth potential.

*Square Grid Model: Enaptation of Robustness* To illustrate the effect created by the force arising from the expected growth potential, Eqs. (8) and (11), consider the model depicted below, panel (a). Each site of a square 12 × 12 grid represents a distinct type, and variations consisting of changes of one type into another correspond to transitions between nearest neighbors.

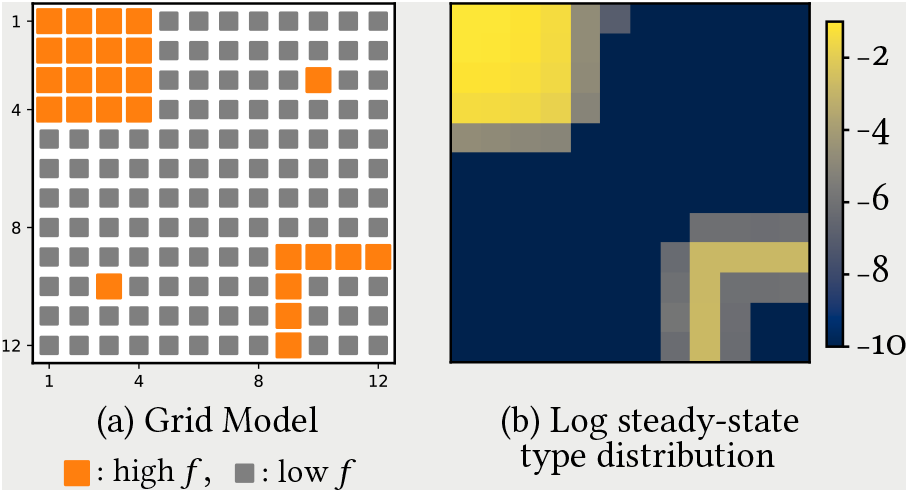

We assume the variation coefficient to be constant and equal to 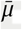. Islands of fast-reproducing types (big orange sites) are surrounded by a sea of slowly reproducing ones (small gray sites), and each island differs by connectivity, *viz*. the number of adjacent slowly growing types. Among types with the same reproduction rate, the expected growth potential favors those surrounded by fast-reproducing types, *viz*. those belonging to the interior of the top-left island. From numerical simulations, the steady-state type distribution (right heat map plot, logarithmic scale) clearly shows a higher probability of finding types surrounded by equally fit types. Indeed, *(i)* the probability of finding an isolated fast-reproducing type (isolated orange sites) is < 10^−10^, and *(ii)* the probability of finding types in the upper square is roughly four order of magnitude higher than that of finding types in the lower one-dimensional strip. This model illustrates how robustness, *i*.*e*. insensitivity to variations, is *enapted* by the evolutionary dynamics.

*Square Grid Model: Fluctuation Relation* To illustrate our fluctuation relation and its implications, we use additional numerical simulations of the grid model. The plot below depicts the average population sensitivity, ⟨*Ω*_*n*_⟩ *vs*. the variation probability log_10_ *μ*. The vertical bars represent one standard deviation, 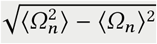.

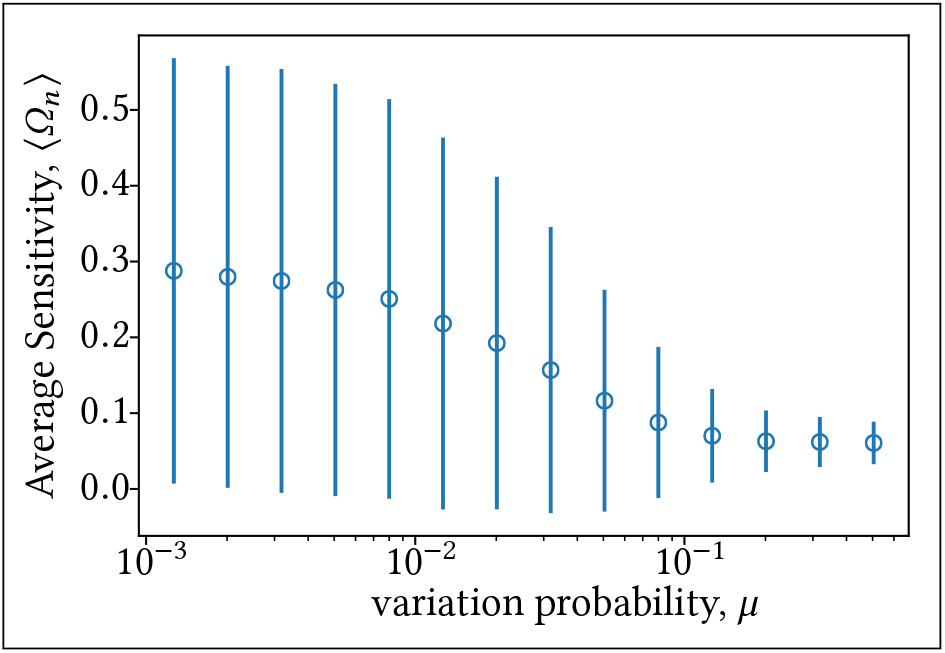

This plot demonstrates the good qualitative agreement between the fluctuation relation and numerical results. As predicted by (15), *(i)* ⟨*Ω*_*n*_⟩ decreases as *μ* increases, and *(ii)* as ⟨*Ω*_*n*_⟩ approaches the plateau, fluctuations decrease as well. We refer to [SI §V] for a more detailed discussion.

### Robustness to variations as an enaptation

Robustness is a concept that has attracted a lot of attention in biology [32–37]. *(i)* Many phenotypic traits of organisms and their developmental pathways have been shown to exhibit robustness to genetic mutations [38], intracellular component concentrations [39], external perturbations [40], etc. There have been also observations of robustness in the behavior of whole multi-species ecological systems [41, 42]. Finally, life on Earth has been until now robust enough to sustain life for billions of years despite strong internal variations and external perturbations. *(ii)* Many studies of model organisms were performed in laboratories to uncover the molecular and cellular mechanisms which underlie robustness [43]. Generic mechanisms—such as redundancy, modularity, and feedback control—have been described in many systems [44–48]. *(iii)* It is difficult to demonstrate, but it is widely believed, that robustness facilitates better survival and reproduction in naturally occurring biological systems [49]. *(iv)* It is possible to show using computer simulations and simple mathematical models that under some important assumptions (*e*.*g*. strong selection, and high mutation rates in sequential genomes) robust variants may outcompete even faster growing variants [50–53].

The general mechanism behind this last phenomenon— which has been referred to as “survival of the flattest”—is elucidated by our analysis. In fact, the existence of a “robustness-generating” evolutionary force points towards a potentially novel mechanism associated with evolution—a mechanism that falls neither under the concept of adaptation nor exaptation [8]. A phenotypic feature, growth robustness, does not emerge in our model as a consequence of selection for robustness because of its fitness value, nor is it co-opted from a previously evolved trait for its new function. Rather, the “robustness-generating” evolutionary force is intrinsic to the evolutionary dynamics itself. This force contribution expresses the tendency of the selected variant to be “surrounded” (in the type space) by other similarly fit variants. We propose to call this class of emergent phenomena *enaptations* (from Latin *en-*, in, into + *aptāre*, to fit), since robustness emerges here as a consequence of intrinsic aspects of evolutionary dynamics *per se*.

We would like to stress again that the emergence of robustness through enaptation does not preclude the existence of other mechanisms which engender this trait, see *e*.*g*. [34].

### Quasispecies, viruses and a fluctuation relation

It might have not escaped the attention of the reader that our model bears some similitude with previous models of evolutionary dynamics like the Wright–Fisher or the *quasispecies* models [15, 54–56], see [SI §VII]. In contrast to these models, we specify neither the nature of *γ*, nor the kind of variations. Quasispecies models belong to the class of our restricted RVS dynamics, since they typically consider genetic types with sequential genomes of finite length, and limit variations to point mutations, whose overall probability is the same for all types. More interestingly, the same assumptions seem to be approximately satisfied for many laboratory experiments studying evolution of virus populations [57, 58]. Indeed, viruses, and in particular RNA viruses, seem well-suited to test the predictions of simple evolutionary models: their growth is fast and can be quantified; their mutation rate is high; and their population size can be easily controlled. It seems, therefore, that we could use the results of such laboratory experiments to at least qualitatively assess the validity of the predictions of our model. To do this it is useful to consider the fluctuation relation derived in our restricted model.

We argued above that under the assumptions of the restricted model, the probability of observing any population composition approximates at long times a Boltzmann-like distribution, Eq. (12). This distribution is ruled by the evolutionary potential Eq. (6) containing the *ϕ*_*n*_ term. The fluctuation relation that ensues, Eq. (15), shows that the robustness to variations of the population increases as a function of the product *μN*, see also Refs. [34, 53]. Most importantly, this relation connects the fluctuations of sensitivity, var{*Ω*_*n*_}, to the changes of the mean sensitivity as a function of the variation rate 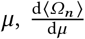. The fluctuation relation further implies that when the sensitivity to variations saturates and approaches a minimum, its fluctuations decrease. Numerical simulations qualitatively verify these predictions, see Box 2 and [SI §V].

Although we derived the enaptation of robustness and the fluctuation relation using the model (5), these results are expected to be approximately valid for any evolutionary stochastic dynamics appropriately including reproduction, variation, and selection (*e*.*g*. the aforementioned quasispecies as well as diffusion models [15, 22]). It is indeed these three elements that are essential here.

### Possible quantitative experiments

Simple predictions of the fluctuation relation seem to be consistent with experimental results involving viral evolution. For instance, in Ref. [59], two sets of viral populations founded from the same lineage were both evolved under strong selection but with different effective mutation rates *μ*. This was achieved by controlling the level of co-infection, which in turn, through complementation, changed the effective probability of mutations *μ*. When subjected to mutation accumulation experiments, viruses evolved under low co-infection (*i*.*e*., under high *μ*) not only showed lower sensitivity to mutations, but also the fluctuations of sensitivity seemed to be significantly lower, see [SI §VIII]. From the perspective of our fluctuation relation, this is a sign that the sensitivity has approached the saturation value. Although this qualitative agreement is encouraging, unfortunately these experimental results do not present us with enough statistics to compare them quantitatively with predictions of evolutionary models. (*E*.*g*. the difference in the mean values of the growth rate between high- and low-mutation rate strains in Ref. [59] is less than one standard deviation of the variations in measurements). However, these interesting experimental results indicate that quantitative comparisons should be possible in principle. Other kinds of experiments which could provide quantitative data to be compared with the predictions of the RSV dynamics are: *(i)* experiments on directed evolution of RNAs [60], proteins [61], or microbial communities [62]; and *(ii)* mutation accumulation experiments involving model organisms [16, 17]. If one is interested in evolution of naturally replicating entities, however, viruses (including bacteriophages) are arguably the best candidates for relevant quantitative experiments.

### Simplistic nature of the model

We conclude with a few additional comments about our model. It is clear that the dynamics described by Eq. (5) is a strong simplification of real evolutionary dynamics. Three major and related aspects have been here neglected: *(i)* For the sake of simplicity, we here disregarded direct interactions among organisms. *De facto*, organisms interact only indirectly through the environment, which affects their reproduction rate, variation probability, and overall population size. We refer to [SI §IB] for the generalization of our model that accounts for interactions. *(ii)* We deliberately focused on constant environments, while real environments do change, fluctuate, and their variations are affected by the population dynamics itself. Although such environmental changes can be formally incorporated in our model, we did not find them relevant at this stage of model development. Environmental changes could be described as an additional stochastic force: it would act alongside the forces described by Eq. (2) and it might favor types that quickly adapt to environmental changes [63–65]. However, such additional forces depend on how the environment alters the population’s reproduction and hence are of a more idiosyncratic nature. Classification of “adaptation strategies” could form a complementary approach to studing the influence of changing environments [66, 67]. *(iii)* We also disregarded any distinctions between genotype, epigenotype and phenotype while introducing variations. This was done for the sake of generality. However, such distinctions definitely become important for changing environments. What differentiates genomes, epigenomes and phenotypes (other than the molecular nature of their realizations) are the characteristic time scales at which they vary. These time scales reflect typical time scales over which information gathered about the environment remains useful. For constant environments considered here, the same information can be used forever, and there is no real need to differentiate between different time scales, and thus between different classes of variations.

In conclusion, we hope that despite its simple nature, our approach may be of general interest. By connecting evolutionary dynamics to non-equilibrium statistical mechanics, it can render more precise some widely used notions, such as “evolutionary forces” or “chance” in evolution.

The authors acknowledge B.K. Xue, P. Sartori, and D. Huse for illuminating discussions. They also thank D. Huse, L. Peliti, M. Esposito, M. Bauer, P. Sartori, N. Lenner, and C.L. Chariker for feedback on the manuscript. The authors are also grateful to M. Hledík for perceptive comments about our model. R.R. was funded by *The Martin A. and Helen Chooljian* Membership in Biology at the Institute for Advanced Study.

