## Supplementary Information for "Evolutionary Dynamics, Evolutionary Forces, and Robustness: A Nonequilibrium Statistical Mechanics Perspective"

### Supplementary Information
### Evolutionary Dynamics, Evolutionary Forces, and Robustness


Riccardo Rao and Stanislas Leibler

*Simons Center for Systems Biology, School of Natural Sciences, Institute for Advanced Study, 08540 Princeton (NJ), U.S.A. and Laboratory of Living Matter, The Rockefeller University, 10065 New York (NY), U.S.A.*



(Dated: October 10, 2021)


The content of these supplementary information is organized as follows. In Sec. I we discuss some technical derivations and generalizations of our Reproduction–Variation–Selection model. In Sec. II, the generalization of our model to continuous variables is introduced. The limiting case of small variation probability is investigated in Sec. III, whereas in in Sec. IV we derive our fluctuation relation. In Secs. V and VI, we discuss the numerical and mathematical details of the toy models introduced in Box 2. The relation between our model and previous ones is discussed in Sec. VII. We conclude by discussing some experiments consistent with our theory, Sec. VIII.

#### I. REPRODUCTION–VARIATION–SELECTION DYNAMICS

In this section we establish our model for reproduction–variation–selection (RVS) dynamics. We start from a generic setting, in which the population size is small and fluctuating, and the organisms composing the population interact, §I A. The limit of large and constant population size is discussed in §I B, and the related evolutionary forces are described in §I C. The model presented in the main text is readily obtained from the aforementioned limit when interactions are neglected. We then move to discuss a few technical aspects that are mentioned in the main text: the exact conditions for a conservative evolutionary dynamics, §I D; and the relation between the expected growth potential and robustness, §I E.

##### A. Fluctuating Population Size

As described in the main text, we consider a population of organisms characterized by some generic hereditary variables $\gamma$. The population composition of a particular generation is denoted by $n = (n_\gamma)$, while its total population by $\mathcal{N}(n) := \sum_\gamma n_\gamma$. In our model, $n$ is a stochastic variable. Let $p_n(\tau)$ describe the probability of observing $n$ at generation $\tau$. At each generation, reproducing organisms are subject to random variations and reproduce. The probability of a $(\gamma \leftarrow \gamma')$-variation is denoted by $\pi_{\gamma\leftarrow\gamma'}(n')$, and the expected reproduction rate of $\gamma$ is denoted by $f_\gamma(n)$. Selection is also random: it acts upon the afore-mentioned variation, and is influenced by the expected reproduction rate of each species, see Fig. 1 in main text. Note that we do not neglect interactions among organisms, and hence both $\pi$ and $f$ may depend on the population composition.

We consider the case for which the population size $\mathcal{N}(n)$ changes due to fluctuations of reproduction and selection. However, we also imagine that the environment keeps the average population constant, which we set to $N$. Mathematically, this dynamics is described by a Chapman–Kolmogorov equation of the form,

$$p_n(\tau+1) = \textstyle\sum_{n'} \mathcal{W}_{n\leftarrow n'} p_{n'}(\tau)\,, \tag{S1}$$

where the transition kernel describes the probability of observing the transition of population composition $n \leftarrow n'$ over one generation. This kernel can be expressed as a product of Poisson processes,

$$\mathcal{W}_{n\leftarrow n'} = \exp\{-N\} \prod_\gamma \frac{1}{n_\gamma!} \left[\Lambda_\gamma(n')\right]^{n_\gamma} \,, \tag{S2}$$

where

$$\Lambda_\gamma(n) := \frac{N f_\gamma(\pi n) \sum_{\gamma'} \pi_{\gamma\leftarrow\gamma'}(n)\, n_{\gamma'}}{\sum_{\gamma'\gamma''} f_{\gamma''}(\pi n)\, \pi_{\gamma''\leftarrow\gamma'}(n)\, n_{\gamma'}} \tag{S3}$$

describes the average growth rate of type $\gamma$ over one generation. In this last expression, one can recognize the stochastic events that characterize one generation: $(\pi n)_\gamma := \sum_{\gamma'} \pi_{\gamma\leftarrow\gamma'}(n)\, n_{\gamma'}$ is the probability that $\gamma$ is engendered by a variation, whereas the reproductive success of this variant is accounted for by $f_\gamma(\pi n)$.

In Eq. (S3), the expected reproduction rate of $\gamma$, $f_\gamma(\pi n) \sum_{\gamma'} \pi_{\gamma\leftarrow\gamma'}(n)\, n_{\gamma'}$ is rescaled so that the average population size remains constant,

$$\sum_n \mathcal{N}(n)\, p_n(\tau+1) = \sum_{nn'} \mathcal{N}(n)\, \mathcal{W}_{n\leftarrow n'} p_{n'}(\tau) = \sum_{\gamma n'} \Lambda_\gamma(n')\, p_{n'}(\tau) = N\,, \quad \text{for all } \tau\,. \tag{S4}$$

where we have used the properties of Poisson distributions. Analogously, the variance of the population size remains constant and scales as $N$,

$$\sigma_{\mathcal{N}}^2 := \sum_n (\mathcal{N}(n) - N)^2 \, p_n(\tau+1) = \sum_{nn'} (\mathcal{N}(n) - N)^2 \, \mathcal{W}_{n\leftarrow n'} p_{n'}(\tau) = \sum_{\gamma n'} \Lambda_\gamma(n') \, p_{n'}(\tau) = N\,, \quad \text{for all } \tau\,. \tag{S5}$$

##### B. Large and Constant Population Size

For large $N$, $N \gg 1$, one finds that $\sigma_{\mathcal{N}}/N = 1/\sqrt{N} \ll 1$. Hence, the fluctuations of population size can be neglected, $\mathcal{N}(n) \simeq N$, and Eq. (S2) can be rewritten as a multinomial,

$$\mathcal{W}_{n\leftarrow n'} \simeq N^N \exp\{-N\} \prod_\gamma \frac{1}{n_\gamma!}\left[\frac{\Lambda_\gamma(n')}{N}\right]^{n_\gamma} \simeq N! \prod_\gamma \frac{1}{n_\gamma!}\left[\frac{f_\gamma(\pi n)\sum_{\gamma'}\pi_{\gamma\leftarrow\gamma'}(n)\, n_{\gamma'}}{\sum_{\gamma'\gamma''} f_{\gamma''}(\pi n)\, \pi_{\gamma''\leftarrow\gamma'}(n)\, n_{\gamma'}}\right]^{n_\gamma}\,. \tag{S6}$$

In the second equality, we have used Stirling approximation and have written explicitly the term in square brackets.

When interactions are neglected, we recover the kernel presented in the main text, Eq. (5).

##### C. Evolutionary Directionality and Evolutionary Forces

As argued in the main text, the log ratio of transition probabilities determines towards the direction towards which population composition is more likely to evolve. This quantity, which we called *evolutionary directionality*, can be generically expressed as the sum of two evolutionary force contributions:

$$\mathcal{F}_{n\leftarrow n'} := \ln \frac{W_{n\leftarrow n'}}{W_{n'\leftarrow n}} = (\psi_n - \psi_{n'}) + \zeta_{nn'}\,. \tag{S7}$$

For the kernel given in Eq. (S6), we obtain

$$\psi_n = \phi_n - \sum_\gamma \ln n_\gamma! \tag{S8}$$

$$\zeta_{nn'} = \ln \prod_\gamma \frac{\left[\sum_{\gamma'} f_\gamma(\pi n')\, \pi_{\gamma\leftarrow\gamma'}(n')\, n'_{\gamma'}\right]^{n_\gamma}}{\left[\sum_{\gamma'} f_\gamma(\pi n)\, \pi_{\gamma\leftarrow\gamma'}(n)\, n_{\gamma'}\right]^{n'_\gamma}}\,, \tag{S9}$$

where

$$\phi_n := N \ln \sum_{\gamma\gamma'} f_\gamma(\pi n)\, \pi_{\gamma\leftarrow\gamma'}(n)\, n_{\gamma'}\,, \tag{S10}$$

denotes the *expected growth potential* (see main text). The approach used to derive Eqs. (S8) and (S9) is to separate the contributions that can be written as a difference of a potential from those that cannot. To do so, we follow a purely algebraic procedure. Since we do not specify the structure of either the type space, or the variations, this procedure is the most general one. In the next subsection, we discuss the special conditions that make the nonconservative contribution vanish.

When neglecting the possibility that interactions affect the reproduction, $f_\gamma(n) = f_\gamma$ for all $n$, then the contribution of selection ($f_\gamma$) to the nonconservative force $\zeta_{nn'}$ becomes conservative and can be regarded as part of the $\psi_n$ term, *cf.* Eqs. (6) and (7) in the main text. The appearance of selection ($f_\gamma$) in the nonconservative force contribution $\zeta_{nn'}$ reflects the more idiosyncratic nature of evolutionary dynamics that involve interactions.

We emphasize that the form of the expected growth potential $\phi_n$ is not affected by the presence of interactions.

##### D. Conditions for Conservative Dynamics

In this section, we analyze under what conditions the dynamics is exactly conservative. This happens when the nonconservative force contribution (S9) can be written as the difference of a potential, and can thus be regarded as an additional contribution of $\psi_n$. As an important consequence, the stationary probability distribution can be written as

$$\bar{p}_n = \frac{\exp \psi_n}{Z}\,, \qquad \text{where} \quad Z := \textstyle\sum_n \exp \psi_n\,. \tag{S11}$$

To identify the conditions for which $\zeta_{nn'}$ can be written as a potential difference (for any $n$ and $n'$) it is crucial to observe that the terms in square brackets (Eq. (S8)) must not depend on the population compositions. In this way, one obtains

$$\zeta_{nn'} = \sum_{\gamma} n_\gamma \ln g_\gamma - \sum_{\gamma} n'_\gamma \ln g_\gamma \,, \tag{S12}$$

for some function of the type $g_\gamma$.

To identify the form of $g_\gamma$ we proceed by steps. First, we need to impose that interactions affect neither the reproduction rate nor the variation probability. Hence,

$$\zeta_{nn'} = \sum_{\gamma} n_\gamma \ln f_\gamma - \sum_{\gamma} n'_\gamma \ln f_\gamma + \ln \prod_{\gamma} \frac{\left[\sum_{\gamma'} \pi_{\gamma\leftarrow\gamma'} n'_{\gamma'}\right]^{n_\gamma}}{\left[\sum_{\gamma'} \pi_{\gamma\leftarrow\gamma'} n_{\gamma'}\right]^{n'_\gamma}} \,, \tag{S13}$$

In other words, the reproduction terms within the nonconservative force contribution (S9) can be written as a potential difference. In the main text (Eqs. (6) and (7)), as well as in the following, these terms are regarded as part of the conservative force described by $\psi_n$ rather than $\zeta_{nn'}$.

Second, the terms in square brackets in Eq. (S13) do not depend on the population composition if and only if the variation probability solely depends on the target type, *i.e.* $\pi_{\gamma\leftarrow\gamma'} = \pi_\gamma$ for all $\gamma'$. In other words, at each generation, each type $\gamma'$ varies into the type $\gamma$ with probability $\pi_\gamma$, irrespectively of $\gamma'$. In this case,

$$\textstyle\sum_{\gamma'} \pi_{\gamma\leftarrow\gamma'} \, n_{\gamma'} = N\pi_\gamma \,, \tag{S14}$$

and one obtains that $g_\gamma = f_\gamma \pi_\gamma$, so that

$$\zeta_{nn'} = \sum_{\gamma} n_\gamma \ln f_\gamma + \sum_{\gamma} n_\gamma \ln \pi_\gamma - \sum_{\gamma} n'_\gamma \ln f_\gamma - \sum_{\gamma} n'_\gamma \ln \pi_\gamma \,. \tag{S15}$$

In this form, it is clear that $\ln \pi_\gamma$ represents how much a certain type is favored by variations.

The potential contributions appearing in (S15) can be thus regarded as a additional contributions to $\psi_n$, Eq. (S8):

$$\psi_n = \sum_{\gamma} n_\gamma \ln f_\gamma + \sum_{\gamma} n_\gamma \ln \pi_\gamma - \sum_{\gamma} \ln n_\gamma! + N \ln \sum_{\gamma} f_\gamma \pi_\gamma \,, \tag{S16}$$

where the last term is the contribution $\phi_n$. The form of $\psi_n$ in Eq. (S16) enters in Eq. (S11).

##### E. Expected Growth Potential and Robustness

As we discuss in the main text, the force contribution corresponding to expected growth potential $\phi_n$ is responsible for driving the evolution of robustness to variations. We now detail the derivation of this result, summarized by Eq. (10) (and (11)) in the main text, and rewritten here for convenience:

$$\phi_n := N \ln \sum_{\gamma\gamma'} f_{\gamma'}(\pi n)\, \pi_{\gamma'\leftarrow\gamma}(n)\, n_\gamma = N \ln \left\{ \textstyle\sum_\gamma f_\gamma(\pi n)\, n_\gamma - \sum_\gamma \mu_\gamma(n)\, \omega_\gamma(n)\, f_\gamma(\pi n)\, n_\gamma \right\} \,, \tag{S17}$$

where

$$\mu_\gamma(n) := \sum_{\gamma'\neq\gamma} \pi_{\gamma'\leftarrow\gamma}(n) \quad \text{and} \tag{S18}$$

$$\omega_\gamma(n) := \sum_{\gamma'} \frac{f_\gamma(\pi n) - f_{\gamma'}(\pi n)}{f_\gamma(\pi n)} \frac{\pi_{\gamma'\leftarrow\gamma}(n)}{\mu_\gamma(n)} \,. \tag{S19}$$

denote the overall variation rate of $\gamma$ and the sensitivity of $\gamma$'s reproduction rate to variations.

To simplify our notation, we omit to report the explicit dependence of $f_\gamma$ and $\pi_{\gamma\leftarrow\gamma'}$ on the actual population composition.

Equation (S17) follows from

$$\begin{aligned}\sum_{\gamma'} f_{\gamma'}\pi_{\gamma'\leftarrow\gamma} &= f_\gamma \pi_{\gamma\leftarrow\gamma} + \sum_{\gamma'\neq\gamma} f_{\gamma'}\pi_{\gamma'\leftarrow\gamma} \\ &= f_\gamma\left[1-\mu_\gamma\right] + \mu_\gamma \sum_{\gamma'\neq\gamma} f_{\gamma'}\frac{\pi_{\gamma'\leftarrow\gamma}}{\mu_\gamma} \\ &= f_\gamma\left\{1-\mu_\gamma\left[1-\sum_{\gamma'\neq\gamma}\frac{f_{\gamma'}}{f_\gamma}\frac{\pi_{\gamma'\leftarrow\gamma}}{\mu_\gamma}\right]\right\} \\ &= f_\gamma\left\{1-\mu_\gamma\sum_{\gamma'\neq\gamma}\left(1-\frac{f_{\gamma'}}{f_\gamma}\right)\frac{\pi_{\gamma'\leftarrow\gamma}}{\mu_\gamma}\right\} \\ &\equiv f_\gamma\left\{1-\mu_\gamma\omega_\gamma\right\}\ .\end{aligned} \tag{S20}$$

To better understand the effects generated by $\phi_n$, let us now consider populations compositions $\bar{n}$ characterized as follows: the reproduction rate of the types in $\bar{n}$ is high compared to their variants and it is roughly constant, $f_\gamma(\pi\bar{n}) \simeq f^*$, for any $\gamma$ such that $\bar{n}_\gamma > 0$. This case could describe situations in which selection is strong enough so that only the fastest-reproducing types can survive.

Hence, Eq. (S17) can be written as

$$\begin{aligned}\phi_{\bar{n}} &= N\ln f_{\text{high}} + N\ln\left\{N - \textstyle\sum_\gamma \mu_\gamma\omega_\gamma\bar{n}_\gamma\right\} \\ &\simeq N\ln\left\{f_{\text{high}}N\right\} - \textstyle\sum_\gamma\mu_\gamma\omega_\gamma\bar{n}_\gamma \\ &\equiv -\textstyle\sum_\gamma\mu_\gamma\omega_\gamma\bar{n}_\gamma + \text{const.}\end{aligned} \tag{S21}$$

The approximation in the second line is justified as the product $\mu_\gamma\omega_\gamma$ is expected to be small: it is the product of two values strictly smaller than one. We thus recover Eq. (11) in the main text. In this form, the force arising from the expected growth potential, $\phi_{\bar{n}} - \phi_{\bar{n}'}$, clearly favors population compositions whose organisms exhibit either low variation probability, low $\mu_\gamma$, or high robustness, low $\omega_\gamma$.

*Multiple variation mechanisms*

As mentioned in the main text, different mechanisms causing variations may be at work simultaneously. Accordingly, the type variation probability can be written as

$$\pi_{\gamma\leftarrow\gamma'} = \textstyle\sum_\nu \pi^{(\nu)}_{\gamma\leftarrow\gamma'}\,, \tag{S22}$$

where $\nu$ labels all mechanisms and their combinations. For instance, if mutations, $\nu = 1$, and recombination, $\nu = 2$, affect a certain organism, then $\nu \in \{1,2,3\}$, where $\nu = 3$ accounts for the probability that a certain organism is subject to both variation mechanisms. Using the equality

$$\mu_\gamma\omega_\gamma = \sum_{\gamma'}\frac{f_\gamma - f_{\gamma'}}{f_\gamma}\pi_{\gamma'\leftarrow\gamma} = \sum_\nu \mu^{(\nu)}_\gamma \sum_{\gamma'}\frac{f_\gamma - f_{\gamma'}}{f_\gamma}\frac{\pi^{(\nu)}_{\gamma'\leftarrow\gamma}}{\mu^{(\nu)}_\gamma} \equiv \sum_\nu \mu^{(\nu)}_\gamma\omega^{(\nu)}_\gamma\,, \tag{S23}$$

Eq. (S17) can be explicitly written in terms of each mechanism

$$\phi_n = N\ln\left\{\textstyle\sum_\gamma f_\gamma n_\gamma - \sum_\nu\sum_\gamma \mu^{(\nu)}_\gamma\omega^{(\nu)}_\gamma f_\gamma n_\gamma\right\}\ . \tag{S24}$$

This implies that the force contribution $\phi_n - \phi_{n'}$ drives the emergence of either robustness, or reduced variation probability, for each single variation mechanism.

*Recombination*

We conclude this subsection by giving an example of variation mechanism involving two organisms: recombination. In this case, the variation probabilities also depend on the population composition,

$$\pi^{(\text{rec})}_{\gamma\leftarrow\gamma'}(n) = \frac{1}{N}\sum_{\gamma''} R_{\gamma\leftarrow\gamma'\gamma''}n_{\gamma''}\,, \tag{S25}$$

where $R_{\gamma\leftarrow\gamma'\gamma''}$ denotes the probability that $\gamma'$ combines with $\gamma''$ and gives birth to $\gamma$. Since $\sum_\gamma R_{\gamma\leftarrow\gamma'\gamma''} = 1$,

$$\mu_{\gamma'}^{(\mathrm{rec})}(n) := \frac{1}{N}\sum_{\gamma\neq\gamma'}\sum_{\gamma''} R_{\gamma\leftarrow\gamma'\gamma''} n_{\gamma''} = 1 - \frac{1}{N}\sum_{\gamma''} R_{\gamma'\leftarrow\gamma'\gamma''} n_{\gamma''} \tag{S26}$$

denotes the probability that any recombination event affects $\gamma$. We can thus express the sensitivity to recombination of $\gamma$ as

$$\omega_{\gamma}^{(\mathrm{rec})}(n) = \frac{1}{N}\sum_{\gamma'\gamma''} \frac{f_\gamma(\pi^{(\mathrm{rec})}n) - f_{\gamma'}(\pi^{(\mathrm{rec})}n)}{f_\gamma(\pi^{(\mathrm{rec})}n)} \frac{R_{\gamma'\leftarrow\gamma\gamma''} n_{\gamma''}}{\mu_\gamma^{(\mathrm{rec})}(n)} \,. \tag{S27}$$

#### II. CONTINUOUS HEREDITARY VARIABLES

In this section, we formulate our model of evolutionary dynamics for continuous hereditary variables. For the sake of simplicity, we disregard interactions between organisms.

Let $\{\,\gamma^{(i)}\,\}$ for $i = 1, \ldots, N$ denote the set of types composing the population. The *population density*, defined as follows, replaces the discrete population composition $n$,

$$\rho(\gamma) := \frac{1}{N}\int \mathrm{d}\gamma \sum_{i=1}^{N} \delta\left(\gamma - \gamma^{(i)}\right) , \tag{S28}$$

where $\delta$ denotes Dirac's deltas, and the integration space spans all possible variables $\gamma$. The probability to observe a certain density $\rho$ at generation $\tau$, $p(\rho; \tau+1)$, is ruled by the equation

$$p(\rho; \tau+1) = \int \mathrm{d}\rho' W(\rho \leftarrow \rho') p(\rho'; \tau) \,. \tag{S29}$$

In a space of continuous hereditary variables, the transition kernel corresponding to Eq. (S6) is given by

$$W(\rho \leftarrow \rho') = N!\, \exp\left\{\int \mathrm{d}\gamma \left[-\ln\left(N\rho(\gamma)\right)! + N\rho(\gamma)\ln\frac{f(\gamma)\int \mathrm{d}\gamma'\bar\pi(\gamma\leftarrow\gamma')\rho(\gamma')}{\int \mathrm{d}\gamma'\mathrm{d}\gamma'' f(\gamma'')\bar\pi(\gamma''\leftarrow\gamma')\rho(\gamma')}\right]\right\} , \tag{S30}$$

where $\bar\pi(\gamma \leftarrow \gamma')$ is the probability density of transitioning from $\gamma'$ to $\gamma$ due to variations.

As in the discrete-space case, the log ratio of forward and backward transition densities determines the evolutionary directionality

$$\mathcal{F}(\rho \leftarrow \rho') := \ln\frac{W(\rho\leftarrow\rho')}{W(\rho'\leftarrow\rho)} = (\psi_\rho - \psi_{\rho'}) + \zeta_{\rho\rho'} \,, \tag{S31}$$

where the force contributions read

$$\psi_\rho := N\int \mathrm{d}\gamma\rho(\gamma)\ln f(\gamma) - \int \mathrm{d}\gamma \ln(N\rho(\gamma))! + N\ln\int \mathrm{d}\gamma\mathrm{d}\gamma' f(\gamma)\bar\pi(\gamma\leftarrow\gamma')\rho(\gamma') \tag{S32}$$

$$\zeta_{\rho\rho'} := N\int \mathrm{d}\gamma\left[\rho(\gamma)\ln\int \mathrm{d}\gamma'\bar\pi(\gamma\leftarrow\gamma')\rho'(\gamma') - \rho'(\gamma)\ln\int \mathrm{d}\gamma'\bar\pi(\gamma\leftarrow\gamma')\rho(\gamma')\right] . \tag{S33}$$

#### III. SMALL VARIATION PROBABILITY REGIME

We here discuss our reproduction–variation–selection dynamics (S6) in the limit of constant and small variation probability, $\mu_\gamma := \sum_{\gamma'(\neq\gamma)} \pi_{\gamma'\leftarrow\gamma} = \mu \ll 1$ for all $\gamma$. We assume no interactions among the organisms of the population. We start by analyzing the transition kernel, and then move to the contributions of the evolutionary directionality.

**Transition Kernel**

In this regime, the transition kernel (S6) (Eq. (1) in the main text) can be approximated as

$$
\begin{aligned}
W_{n\leftarrow n'} &= N!\prod_{\gamma}\frac{1}{n_\gamma!}\left[\frac{f_\gamma\left[(1-\mu)\,n'_\gamma+\mu\sum_{\gamma'(\neq\gamma)}\bar{\pi}_{\gamma\leftarrow\gamma'}n'_{\gamma'}\right]}{\sum_{\gamma''}f_{\gamma''}\left[(1-\mu)\,n'_{\gamma''}+\mu\sum_{\gamma'(\neq\gamma'')}\bar{\pi}_{\gamma''\leftarrow\gamma'}n'_{\gamma'}\right]}\right]^{n_\gamma}\\
&\simeq N!\left\{\prod_{\gamma:n'_\gamma\neq0}\frac{1}{n_\gamma!}\left[\frac{f_\gamma n'_\gamma}{\sum_{\gamma'}f_{\gamma'}n'_{\gamma'}}\right]^{n_\gamma}\right\}\left\{\prod_{\gamma:n'_\gamma=0}\frac{1}{n_\gamma!}\left[\frac{\mu f_\gamma\sum_{\gamma'(\neq\gamma)}\bar{\pi}_{\gamma\leftarrow\gamma'}n'_{\gamma'}}{\sum_{\gamma'}f_{\gamma'}n'_{\gamma'}}\right]^{n_\gamma}\right\}\\
&= \mu^{\sum_{\gamma:n'_\gamma=0}n_\gamma}N!\left\{\prod_{\gamma:n'_\gamma\neq0}\frac{1}{n_\gamma!}\left[\frac{f_\gamma n'_\gamma}{\sum_{\gamma'}f_{\gamma'}n'_{\gamma'}}\right]^{n_\gamma}\right\}\left\{\prod_{\gamma:n'_\gamma=0}\frac{1}{n_\gamma!}\left[\frac{f_\gamma\sum_{\gamma'(\neq\gamma)}\bar{\pi}_{\gamma\leftarrow\gamma'}n'_{\gamma'}}{\sum_{\gamma'}f_{\gamma'}n'_{\gamma'}}\right]^{n_\gamma}\right\},
\end{aligned}
\tag{S34}
$$

where we introduced the conditional probability of transitioning

$$
\bar{\pi}_{\gamma\leftarrow\gamma'}=\pi_{\gamma\leftarrow\gamma'}/\mu\,,\quad \text{for } \gamma\neq\gamma'\,,
\tag{S35}
$$

and used the following equality in the first line

$$
\sum_{\gamma'}\pi_{\gamma\leftarrow\gamma'}n_{\gamma'}=(1-\mu)\,n_\gamma+\mu\sum_{\gamma'(\neq\gamma)}\frac{\pi_{\gamma\leftarrow\gamma'}}{\mu}n_{\gamma'}=(1-\mu)\,n_\gamma+\mu\sum_{\gamma'(\neq\gamma)}\bar{\pi}_{\gamma\leftarrow\gamma'}n_{\gamma'}\,.
\tag{S36}
$$

Importantly, we regard $\bar{\pi}_{\gamma\leftarrow\gamma'}$ as independent on $\mu$. In the second line of Eq. (S34), the first term in curly brackets accounts for the selection of types that are present in the population $n'$. This term is dominated by selection and selection fluctuations, since it accounts for the selection of types that either grow faster (higher $f_\gamma$) or are more abundant (higher $n_\gamma$). The second term accounts for the selection of types that appear due to variations, and is strongly dependent on the probabilities of variation. As highlighted in the third line, $W_{n\leftarrow n'}$ scales as a power of $\mu$, where the exponent is the number of new types appearing in the population.

For sufficiently small variation probability, $\mu\ll1/N$, it is unlikely that more than one variation occurs in one generation. Therefore, the typical transition $n\leftarrow n'$ involves either no variation,

$$
W_{n\leftarrow n'}=N!\prod_{\gamma:n'_\gamma\neq0}\frac{1}{n_\gamma!}\left[\frac{f_\gamma n'_\gamma}{\sum_{\gamma'}f_{\gamma'}n'_{\gamma'}}\right]^{n_\gamma},\quad \text{and } n_\gamma=0 \text{ if } n'_\gamma=0, \text{ for all } \gamma\,,
\tag{S37}
$$

or just one,

$$
W_{n\leftarrow n'}=\mu N!\left\{\prod_{\gamma:n'_\gamma\neq0}\frac{1}{n_\gamma!}\left[\frac{f_\gamma n'_\gamma}{\sum_{\gamma'}f_{\gamma'}n'_{\gamma'}}\right]^{n_\gamma}\right\}\frac{f_{\bar{\gamma}}\sum_{\gamma'(\neq\bar{\gamma})}\bar{\pi}_{\bar{\gamma}\leftarrow\gamma'}n'_{\gamma'}}{\sum_{\gamma'}f_{\gamma'}n'_{\gamma'}}\,,
\tag{S38}
$$

where $n_\gamma=0$ if $n'_\gamma=0$, for all $\gamma$ except for the type originating from the variation, $\bar{\gamma}$, for which $n_{\bar{\gamma}}=1$ and $n'_{\bar{\gamma}}=0$.

To fix the ideas, imagine a population composed of just one type $\gamma$ that transitions toward a composition where one new type, $\bar{\gamma}$, appears. The probability of such transition can be written in simple terms,

$$
W_{n\leftarrow n'}=\mu N f_{\bar{\gamma}}\bar{\pi}_{\bar{\gamma}\leftarrow\gamma}/f_\gamma\,.
\tag{S39}
$$

In the limit of large population size, we can also express the probability of the reverse transition in simple terms,

$$
W_{n'\leftarrow n}=\left[\frac{(N-1)\,f_\gamma}{f_{\bar{\gamma}}+(N-1)\,f_\gamma}\right]^N\simeq\exp\left\{-\frac{f_{\bar{\gamma}}}{f_\gamma}\right\}\,.
\tag{S40}
$$

**Evolutionary Directionality**

We now turn our attention to the force contributions of the evolutionary directionality for small $\mu$, Eqs. (6) and (7) in the main text.

*Conservative Force*

By Taylor expanding the variants growth potential $\phi_n$ in terms of $\mu$, we obtain

$$\phi_n := N\ln\sum_{\gamma\gamma'} f_{\gamma'}\pi_{\gamma'\leftarrow\gamma}n_\gamma = N\ln\left\{\sum_\gamma f_\gamma n_\gamma - \mu\left[\sum_\gamma f_\gamma n_\gamma - \sum_{\gamma,\gamma'(\neq\gamma)} f_{\gamma'}\frac{\pi_{\gamma'\leftarrow\gamma}}{\mu}n_\gamma\right]\right\} \tag{S41}$$

$$\simeq N\ln\sum_\gamma f_\gamma n_\gamma - \mu N\frac{\sum_\gamma f_\gamma n_\gamma - \sum_{\gamma,\gamma'(\neq\gamma)} f_{\gamma'}\pi_{\gamma'\leftarrow\gamma}n_\gamma/\mu}{\sum_\gamma f_\gamma n_\gamma} \tag{S42}$$

$$\equiv N\ln F_n - \mu N\frac{F_n - F_{\pi n}}{F_n}\,, \tag{S43}$$

where the first equality follows from

$$\sum_{\gamma'} f_{\gamma'}\pi_{\gamma'\leftarrow\gamma} = f_\gamma\pi_{\gamma\leftarrow\gamma} + \sum_{\gamma'(\neq\gamma)} f_{\gamma'}\pi_{\gamma'\leftarrow\gamma} = f_\gamma(1-\mu) + \mu\sum_{\gamma'(\neq\gamma)} f_{\gamma'}\frac{\pi_{\gamma'\leftarrow\gamma}}{\mu} = f_\gamma - \mu\left[f_\gamma - \sum_{\gamma'(\neq\gamma)} f_{\gamma'}\frac{\pi_{\gamma'\leftarrow\gamma}}{\mu}\right]. \tag{S44}$$

In Eq. (S43), we have introduced the cumulative growth of the population, $F_n := \sum_\gamma f_\gamma n_\gamma$, and the cumulative growth after variations, $F_{\pi n} := \sum_{\gamma,\gamma'(\neq\gamma)} f_{\gamma'}\left(\pi_{\gamma'\leftarrow\gamma}/\mu\right)n_\gamma$. The latter explicitly depends on the conditional probability of variation $\pi_{\gamma'\leftarrow\gamma}/\mu$, for $\gamma'\neq\gamma$. The conservative force potential is therefore

$$\psi_n \simeq \textstyle\sum_\gamma n_\gamma\ln f_\gamma - \sum_\gamma \ln n_\gamma! + N\ln\sum_\gamma n_\gamma f_\gamma - \mu N\dfrac{F_n - F_{\pi n}}{F_n}\,. \tag{S45}$$

Hence, the zeroth contribution—*i.e.* the first three terms on the r.h.s.—accounts for the selection and entropic contributions, and favors populations with higher reproduction and higher diversity. We will see later, §IV, that the first order contribution of the expansion in $\mu$—*i.e.* the last term—favors robust populations.

*Nonconservative Force*

To characterize the nonconservative contribution (S13) we preliminary rewrite it as

$$\zeta_{nn'} := \sum_\gamma \ln\frac{\left[(1-\mu)n'_\gamma + \mu\sum_{\gamma'(\neq\gamma)}\bar\pi_{\gamma\leftarrow\gamma'}n'_{\gamma'}\right]^{n_\gamma}}{\left[(1-\mu)n_\gamma + \mu\sum_{\gamma'(\neq\gamma)}\bar\pi_{\gamma\leftarrow\gamma'}n_{\gamma'}\right]^{n'_\gamma}}\,. \tag{S46}$$

where we used Eq. (S36). We now approximate each logarithmic term in the sum for $\mu\simeq 0$,

$$\ln\frac{\left[(1-\mu)n'_\gamma + \mu\sum_{\gamma'(\neq\gamma)}\bar\pi_{\gamma\leftarrow\gamma'}n'_{\gamma'}\right]^{n_\gamma}}{\left[(1-\mu)n_\gamma + \mu\sum_{\gamma'(\neq\gamma)}\bar\pi_{\gamma\leftarrow\gamma'}n_{\gamma'}\right]^{n'_\gamma}} \simeq \begin{cases} n_\gamma\ln n'_\gamma - n'_\gamma\ln n_\gamma & \text{if } n_\gamma\neq 0, n'_\gamma\neq 0\\ n_\gamma\ln\mu + n_\gamma\ln\sum_{\gamma'}\bar\pi_{\gamma\leftarrow\gamma'}n'_{\gamma'} & \text{if } n_\gamma\neq 0, n'_\gamma = 0\\ -n'_\gamma\ln\mu - n'_\gamma\ln\sum_{\gamma'}\bar\pi_{\gamma\leftarrow\gamma'}n_{\gamma'} & \text{if } n_\gamma = 0, n'_\gamma\neq 0\end{cases} \tag{S47}$$

We can rewrite (S46) as

$$\zeta_{nn'} \simeq -\ell_{nn'}\ln\mu + \sum_{\substack{\gamma:n_\gamma\neq 0,\\ n'_\gamma\neq 0}}\left[n_\gamma\ln n'_\gamma - n'_\gamma\ln n_\gamma\right] + \sum_{\gamma:n'_\gamma=0} n_\gamma\ln\textstyle\sum_{\gamma'}\bar\pi_{\gamma\leftarrow\gamma'}n'_{\gamma'} - \displaystyle\sum_{\gamma:n_\gamma=0} n'_\gamma\ln\textstyle\sum_{\gamma'}\bar\pi_{\gamma\leftarrow\gamma'}n_{\gamma'}\,, \tag{S48}$$

where the coefficient appearing in the first term on the right hand side,

$$\ell_{nn'} := \sum_{\gamma:n_\gamma=0} n'_\gamma - \sum_{\gamma:n'_\gamma=0} n_\gamma\,, \tag{S49}$$

can be interpreted as the net number of types lost in the transition $n \leftarrow n'$. Indeed, $\sum_{\gamma: n_\gamma = 0} n'_\gamma$ quantifies the number of types that were present in $n'$ but disappear in $n$. Vice versa, $\sum_{\gamma: n'_\gamma = 0} n_\gamma$ is the number of types that are present in $n$ but were absent in $n'$.

For $\ell_{nn'} \neq 0$, the force $\zeta_{nn'}$ diverges logarithmically in the limit $\mu \longrightarrow 0$, and the sign of $\ell_{nn'}$ determines the sign of the force. Specifically, $\zeta_{nn'}$ is positive when the net loss of types is larger in the direction $n \leftarrow n'$. This result is in agreement with the observation that selection fluctuations disfavor diversity since the selection of the most abundant types is privileged, see *e.g.* Ref. [1].

It is important to observe that this effect is partly balanced the entropic term in Eq. (S8), which instead favors diversity. By using Stirling approximation, we can write the sum of the entropic and nonconservative force as,

$$\zeta_{nn'} - \sum_\gamma \ln \frac{n_\gamma!}{n'_\gamma!} \simeq \sum_{\substack{\gamma: n'_\gamma = 0, \\ n_\gamma \neq 0}} n_\gamma \ln \frac{\mu N}{n_\gamma} - \sum_{\substack{\gamma: n'_\gamma \neq 0, \\ n_\gamma = 0}} n'_\gamma \ln \frac{\mu N}{n'_\gamma} - \sum_{\substack{\gamma: n'_\gamma \neq 0, \\ n_\gamma \neq 0}} \left[ n_\gamma \ln \frac{n_\gamma}{n'_\gamma} - n'_\gamma \ln \frac{n'_\gamma}{n_\gamma} \right] \tag{S50}$$

where we also neglected the last two terms in (S48). The major contribution of this expression comes again from the first two terms. Indeed, since variations are rare, typical transitions do not involve the appearance of many individuals of the same type, *viz.* if $n'_\gamma = 0$ and $n_\gamma \neq 0$ then $n_\gamma \simeq 1$. Analogously, if $n'_\gamma \neq 0$ and $n_\gamma = 0$ then $n'_\gamma \simeq 1$. Also, typical transitions keep the composition of the abundant types roughly constant, $n_\gamma \sim n'_\gamma$, which implies that the last term in Eq. (S50) is roughly zero. Therefore,

$$\zeta_{nn'} - \sum_\gamma \ln \frac{n_\gamma!}{n'_\gamma!} \simeq -\ell_{nn'} \ln \mu N \, . \tag{S51}$$

This expression highlights how type diversity is preserved or disfavored, depending on whether $\mu N$ is greater or smaller than one. In particular, for $\mu N \ll 1$, the nonconservative force strongly prevails over the entropic one, and diversity is disfavored.

As an illustration, we consider again the case of a transition in which one type $\bar{\gamma}$ appears from a monomorphic population of type $\gamma$ for $\mu N \ll 1$. In this special case, the force contributions are

$$\psi_n - \psi_{n'} \simeq \ln \frac{f_{\bar{\gamma}}}{f_\gamma} + \ln N + \frac{f_{\bar{\gamma}}}{f_\gamma} \tag{S52}$$

$$\zeta_{nn'} = \ln \mu + \ln \bar{\pi}_{\bar{\gamma} \leftarrow \gamma} \, , \tag{S53}$$

where we neglected the first-order contribution in $\mu$ in $\psi_n$. Equations (S53) are in agreement with the result obtained by taking the log ratio of Eqs. (S39) and (S40). For variations among types with equal growth, $f_{\bar{\gamma}}/f_\gamma \simeq 1$, the above expression clarifies that the appearance of a new type is *(i)* driven by the $\ln N > 0$ term coming from the entropic contribution, and *(ii)* prevented by the $\ln \mu < 0$ term coming from the nonconservative one.

#### IV. FLUCTUATION RELATION

In this section, we detail the derivation of the fluctuation relation, Eq. (14) in the main text. To do so, we first discuss a formal definition of strength of selection and strong selection regime.

##### A. Strong Selection Regime

In the strong selection regime, only those types with a high reproduction rate survive. This regime can be formalized by rewriting the reproduction rates as

$$f_\gamma = \exp\left\{\beta \lambda_\gamma\right\} \, . \tag{S54}$$

The whole exponent can be interpreted as the logarithmic growth rate parametrized by $\beta$. For very large $\beta$, tiny differences of $\lambda_\gamma$ give huge differences in $f_\gamma$, and thus types are highly discriminated. In contrast, for small $\beta$, even very large differences in $\lambda_\gamma$ give small differences in $f_\gamma$. Hence, $\beta$ captures the strength of selection: $\beta \gg 1$ implies strong selection.

##### B. Derivation of the Fluctuation Relation

To derive our fluctuation relation, Eq. (15) in main text, we first consider a regime in which the variation probabilities $\mu$s are not too small and selection is strong. Hence, the nonconservative force $\zeta_{nn'}$ remains finite (see Sec. III) and can be regarded as unimportant compared to the conservative one, $(\psi_n - \psi_{n'})$, since $\zeta_{nn'}$ does not account for selection. Hence, we neglect the $\zeta_{nn'}$.

Under this assumption, the dynamics is approximately conservative and the probability distribution $p_n(\tau)$ described by the master equation (Eq. (1) in the main text) relaxes to a Boltzmann–like distribution

$$p_n(\infty) \simeq \frac{\exp \psi_n}{Z}\,, \qquad \text{where} \qquad Z := \textstyle\sum_n \exp \psi_n\,. \tag{S55}$$

where $\psi_n$ is given in Eq. (S60).

Second, we recall the expression of the conservative force potential expanded in $\mu$, Eq. (S43), rewritten here for convenience,

$$\phi_n \equiv N \ln F_n - \mu N \frac{F_n - F_{\pi n}}{F_n}\,. \tag{S56}$$

with

$$F_n := \sum_\gamma f_\gamma n_\gamma \tag{S57}$$

$$F_{\pi n} := \sum_{\gamma,\gamma'(\neq\gamma)} f_{\gamma'} \pi_{\gamma'\leftarrow\gamma} n_\gamma / \mu \tag{S58}$$

being the cumulative reproduction rate prior and after variations, respectively. In analogy to the sensitivity introduced in Eq. (S19), the last term of Eq. (S56) can be interpreted as the sensitivity to variations of the population growth,

$$\Omega_n := \frac{F_n - F_{\pi n}}{F_n}\,. \tag{S59}$$

Hence, the whole potential $\psi_n$ can be rewritten as

$$\psi_n \simeq \textstyle\sum_\gamma n_\gamma \ln f_\gamma - \sum_\gamma \ln n_\gamma! + N \ln F_n - \mu N \Omega_n\,. \tag{S60}$$

Our sensitivity–fluctuation relation then follows from straightforward calculations:

$$\frac{1}{N}\frac{\mathrm{d} \ln Z}{\mathrm{d}\mu} \simeq -\langle \Omega_n \rangle \tag{S61}$$

and

$$-\frac{1}{N}\frac{\mathrm{d}\langle \Omega_n \rangle}{\mathrm{d}\mu} \simeq \frac{1}{N^2}\frac{\mathrm{d}^2 \ln Z}{\mathrm{d}\mu^2} \simeq \langle \Omega_n^2 \rangle - \langle \Omega_n \rangle^2\,. \tag{S62}$$

We recall that the averages read $\langle \Omega_n \rangle = \sum_n p_n(\infty)\Omega_n$.

*Remark* The fluctuation relation bears some similitude with Fisher's fundamental relation, see *e.g.* [2]. To clarify this point, we remark that the latter relation equates the change in time of the mean reproduction rate to the variance of reproduction rate, and is exactly valid only in absence of variations. Our fluctuation relation, instead, is an approximate statement about the sensitivity of reproduction to variations at steady state, rather than along the transient evolution. Hence, we conclude that the similitude is purely mathematical.

#### V. NUMERICAL RESULTS

We here report a numerical test of the fluctuation relation (S62). To do so, we first summarize the evolutionary algorithm and the model that we use.

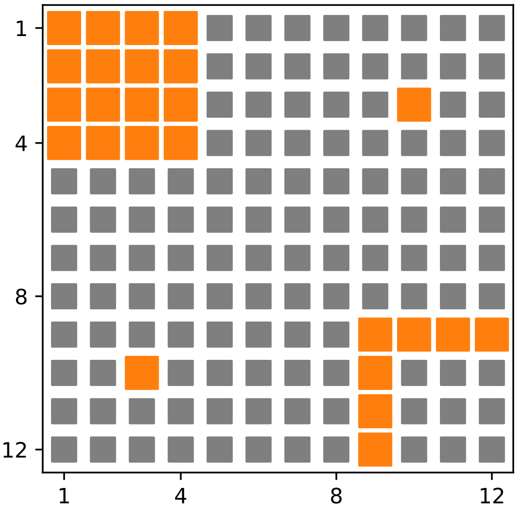


FIG. 1. Type space and values of the log growth function for our grid model. Each square represents a possible type. Large orange squares denote fast growing types $\lambda_\gamma = \lambda_* = 5$, Eq. (S54), whereas small gray squares denote slowly growing ones, $\lambda_\gamma = \lambda_- = 1$.

##### A. Evolutionary Algorithms

Evolutionary algorithms simulate evolutionary dynamics for the purpose of optimization. The simplest version of this class of algorithms can be summarized as follows (for a general introduction to evolutionary optimization algorithms see *e.g.* Ref. [3]).

One starts with a population of $N$ organisms, each of which is characterized by some hereditary variables $\gamma$. Each iteration then consists of the following steps:

1. The hereditary variables of each organism are randomly varied according to the probability $\pi_{\gamma' \leftarrow \gamma}$. The outcome of this step is thus a population of variants, each of which is characterized by a certain reproduction rate $f_\gamma$.

2. $N$ new organisms are sampled with replacement using the reproduction rate of the mutants as a measure. The sampled population goes on to the next iteration.

##### B. Grid Model

We consider a two-dimensional $12 \times 12$ square grid in which each site represents a possible type, Fig. 1. Variations of one type into another correspond to nearest neighbors transitions, and we set the variation coefficient to a constant $\bar{\mu}$. Hence, the overall variation probability is $4\bar{\mu}$ for the types in the bulk, and $3\bar{\mu}$ or $2\bar{\mu}$ for the types at the edges.

The reproduction rate is parametrized as in Eq. (S54), and the values of the function $\lambda_\gamma$ are depicted in Fig. 1. Islands of fast growing types (large orange squares, $\lambda_\gamma = \lambda_* = 5$) are surrounded by a sea of slow ones (small gray squares, $\lambda_\gamma = \lambda_- = 1$), and each island differs by topology, *i.e.* the number of adjacent slowly growing types. In this way, the distribution of fast reproducing types can be effectively probed for different parameters values.

Figure 2, shows the type distributions obtained for different values of the strength of selection $\beta$ and the variation coefficient $\bar{\mu}$. It is worth emphasizing a few observations.

First, we notice the increase of robustness when the distribution of fast reproducing types is heterogeneous. Types belonging to the fast-reproducing two-dimensional island (top left corner) have a higher probability density than those located on the one- and zero-dimensional islands. Even within the two-dimensional island, internal types, which are completely surrounded by fast-growing types, are favored against external ones. Second, except for extremely high strengths of selection and extremely low variation coefficients (top left corner), slowly-reproducing neighbors of the fast-reproducing types are systematically observed, $p_\gamma \gg 1/N_{\text{samples}}(= 10^{-10})$. Both of these two effects become more and more pronounced as the variation coefficient $\bar{\mu}$ is increased.

These observations are in agreement with the phenomenon known as *survival of the flattest*, which originated in the context of the quasispecies theory (see Sec. VII A and Refs. [4, 5]) and can be summarized as follows. For high mutation rates, genotypes that occupy flatter regions of the fitness landscape may have higher chances to survive than types that occupy fitter but narrower regions. This is because mutations are more likely to harm the latter kind of types.

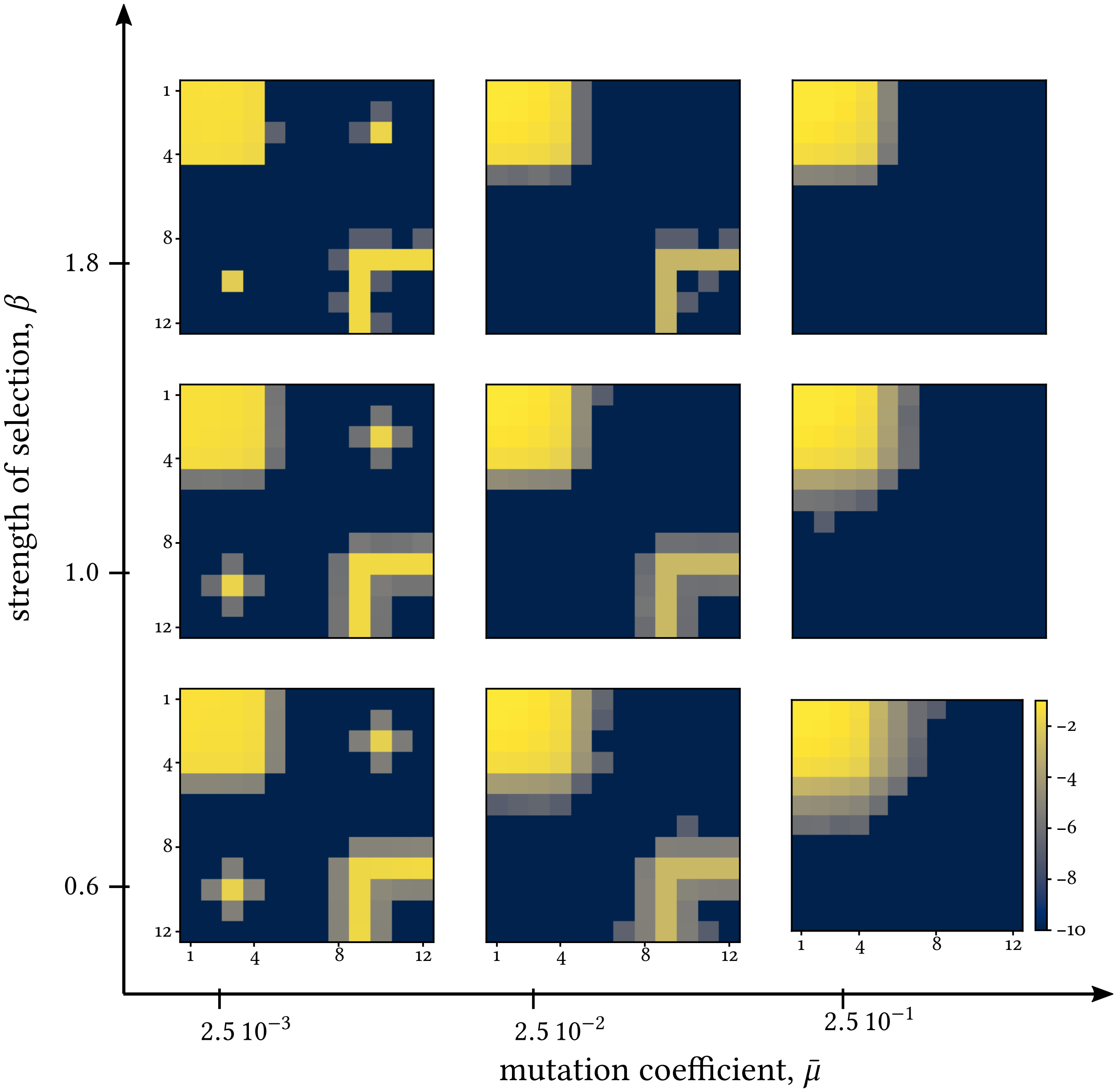


FIG. 2. Log of long-time type probability distributions, $\log_{10} p_Y(\infty)$, obtained for different values of $\beta$ and $\bar{\mu}$. Each distribution is obtained by simulating $10^6$ independent populations of $N = 100$ organisms. For each population, we let the dynamics relax for $10^4$ generations, and then sampled the type of one individual for any of the remaining $10^4$ generations.

Note however that our model generalizes the phenomenon of *survival of the flattest* to dynamics involving generic types and all kinds of variation. It also provides a mechanistic understanding of this phenomenon. Variations and selection generate an effective interaction among the organisms composing the population. This interaction is captured by the expected growth potential $\phi_n$, Eq. (S17). As discussed in the main text, $\phi_n$ favors types surrounded by fast-growing ones and, for type-independent variation probability $\mu$, this effect is stronger at higher $\mu$.

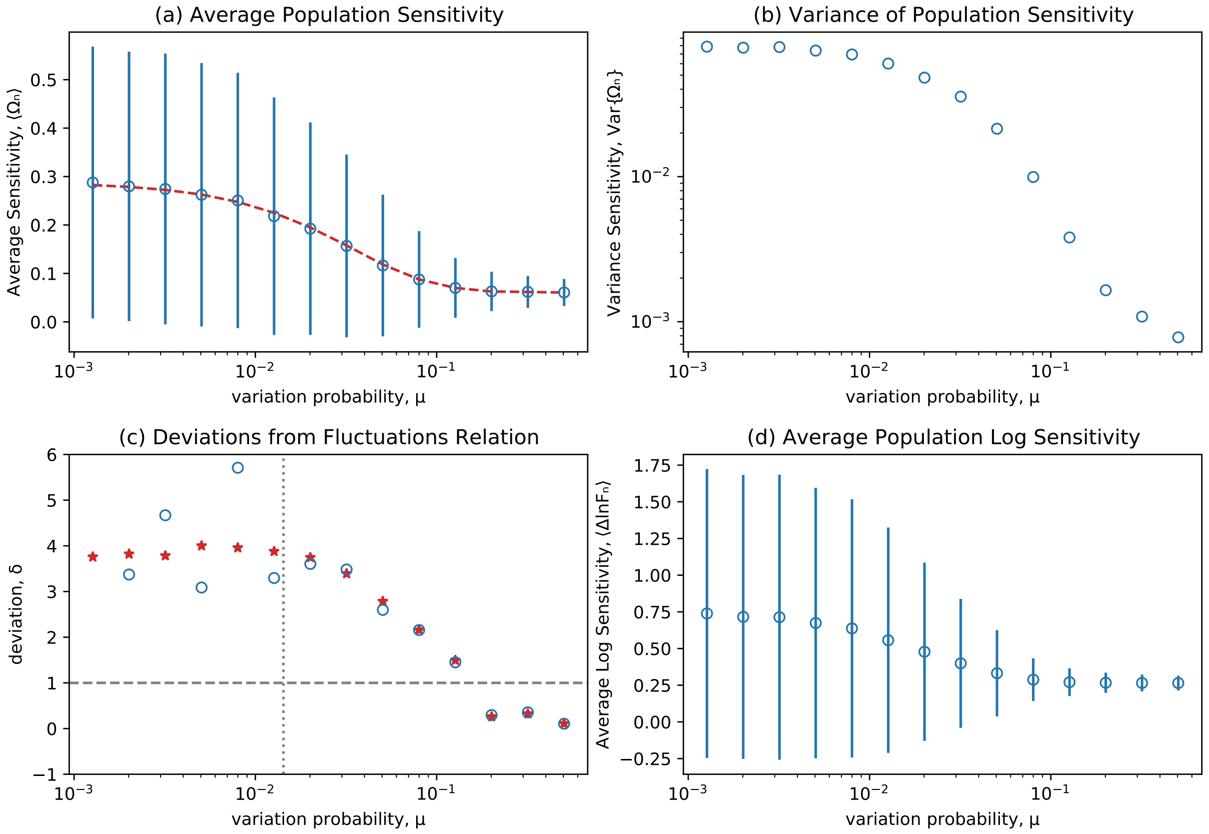


FIG. 3. Numerical analysis of the fluctuation relation. (a) Linear-log plot of the average population sensitivity vs the variation probability $\mu$ (blue circles). The vertical bars denote one standard deviation away from the mean. The red dashed line represents a smoothing B-spline of degree 1 obtained from the data (smoothing factor = $10^{-4}$). (b) Logarithmic plot of the variance of population sensitivity vs the variation probability $\bar{\mu}$. (c) Deviations from fluctuation relation (S62) as quantified by Eq. (S63). Blue circles represent deviations computed using finite difference of $\langle \Omega_n \rangle$ *in lieu* of derivatives, whereas red stars represent deviations computed using the smoothing spline described in figure a. The latter approach is required since the fluctuations of $\langle \Omega_n \rangle$ become very intense at low values of $\mu$. The horizontal gray dashed line ($\delta = 1$) denotes the values for which the fluctuation relation is perfectly satisfied. The vertical gray dotted line represents $\mu = 1/N$. (d) Linear-log plot of average population log sensitivity (Eq. (S64)) vs the variation probability $\mu$. Each data point is estimated from $8 \cdot 10^3$ randomly sampled populations with $N = 70$ and $\beta = 1$.

##### C. Sensitivity and Fluctuation Relation

To systematically quantify the population robustness and test the fluctuation relation, we now focus on $\beta = 1$. Figure 3 plots the statistics $\Omega_n$ at different values of the variation probability $\mu$. Since not all types vary with the same probability (see earlier discussion about types at the boundary) we set the variation probability as $\mu = 4\bar{\mu}$. This assumption does not significantly affect the results of our simulations. At the same time, it shows that our fluctuation relation is robust to cases in which $\mu$ is slightly different for different types.

Figure 3 demonstrates the good qualitative agreement between our fluctuation relation and numerical results. The average population sensitivity, $\langle \Omega_n \rangle$, is plotted against $\log_{10} \mu$ in Fig. 3a, where the vertical bars represent one standard deviation. As predicted by (S62), *(i)* $\langle \Omega_n \rangle$ decreases with $\mu$, and *(ii)* as $\langle \Omega_n \rangle$ approaches the plateau, fluctuations decrease as well. $\mathrm{var}\,\{\Omega_n\} := \langle \Omega^2 \rangle - \langle \Omega \rangle^2$ is plotted vs $\log_{10} \mu$ on a logarithmic scale in Fig. 3b.

Figure 3c plots deviations from the fluctuation relation,

$$\delta := -\frac{\frac{1}{N}\frac{\mathrm{d}\langle \Omega_n \rangle}{\mathrm{d}\mu}}{\langle \Omega^2 \rangle - \langle \Omega \rangle^2}\,, \tag{S63}$$

as function of $\log_{10} \mu$. Overall we find a qualitatively good agreement as $\delta$ is of the order of 1 ($\delta = 1$ corresponds to perfect agreement). At the same time, we confirm that the largest deviations are found for small $\mu$. Indeed, we recall that, for some $n$ and $n'$, the nonconservative force $\zeta_{nn'}$ diverges logarithmically for $\mu \to 0$. This makes the approximation (S55) not fully valid.

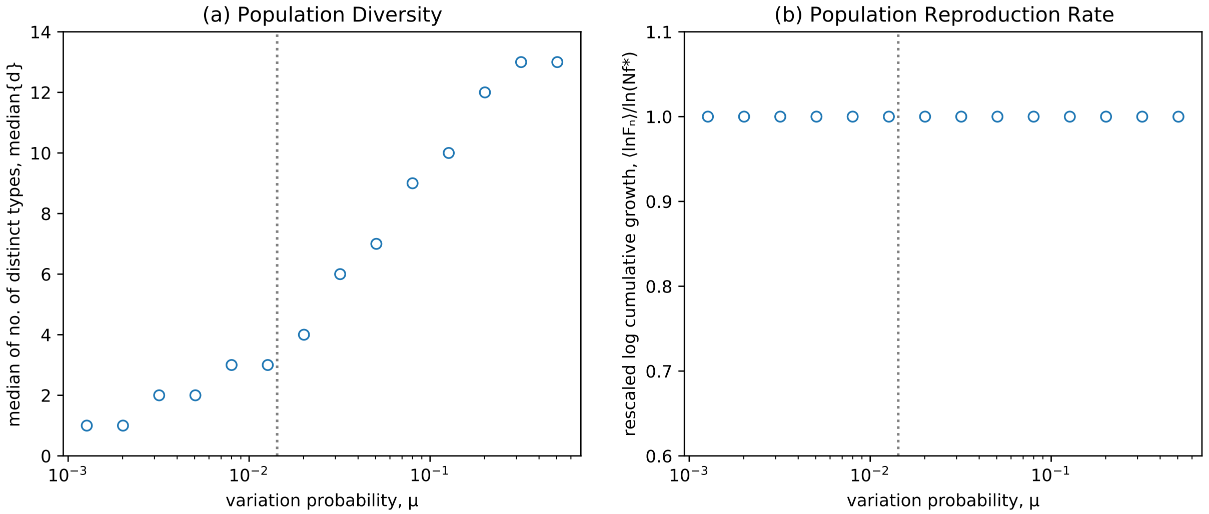


FIG. 4. (a) Median of the number of different types found in the population. The data points are estimated form the same simulation as in Fig. 3. The gray dotted vertical line represents $\mu = 1/N$. For $\mu < 1/N$ no more than 3 distinct types (over 25 highly growing types) typically compose the population. This is in agreement with the average number of distinct types expected from a heuristic argument based on the coefficient $\mu N$: for $\mu N \ll 1$, few variations arise and the population is typically composed of just a few types; for $\mu N \gg 1$, many variations arise, and the population is composed by many distinct types. (b) Rescaled log cumulative growth of the population, $\langle \ln F_n \rangle / \ln(f^* N)$. The almost constant value of 1 indicates that the population is always characterized by a maximal growth.

The fact that such deviations saturate (rather than diverge), can be explained as follows. In the small $\mu$ regime, *(i)* variations are very rare, and *(ii)* the population becomes frozen into configurations characterized by just a few distinct types equally abundant, see Fig. 4(a). Hence, pairs of population compositions for which one type appears or disappears become rare. This means that the population compositions for which $\zeta_{nn'}$ diverges rarely appear, and such force does not completely influence the dynamics.

We finally check our result using a different definition of sensitivity,

$$\Delta \ln F_n := \ln \frac{F_n}{F_{\pi n}}, \tag{S64}$$

which is the one typically used in evolutionary studies of viruses, see *e.g.* [6]. Figure 3d shows this average log sensitivity as function of $\log \mu$. It is clear that no significant difference with respect to Fig. 3a appears. Mathematically, this can be explained by the fact that, for low values of sensitivity, $F_n/F_{\pi n} \simeq 1$, the two definitions are almost equivalent,

$$\Delta \ln F_n \equiv \ln \frac{F_n}{F_{\pi n}} \simeq \frac{F_{\pi n}}{F_n} \left( \frac{F_n}{F_{\pi n}} - 1 \right) \equiv \Omega_n . \tag{S65}$$

*Remark* The behavior of the population for different values of $\mu$ depicted in Fig. 3(a) can also be used to further strengthen the relation between the emergence of robustness and the force arising from the variant growth potential $\phi_n$. To see this, we could consider the "work" done by this force along a transformation for which robustness increases. If this "work" is positive, then this force actively contributes to generate robustness. The transformation that we consider is characterized by a constant and high $\mu$, let us say $10^{-1}$, but the population is initially distributed as for a low $\mu$, let us say $10^{-3}$. In other words, even though $\mu$ is high, the initial population has a high average sensitivity (low robustness), $\langle \Omega_n \rangle_{\mathrm{i}} \simeq 0.3$, Fig. 3(a). As the population relaxes to the steady-state distribution, the sensitivity reaches its minimum $\langle \Omega_n \rangle_{\mathrm{f}} \simeq 0.1$. Since the force originating from $\phi_n$ is conservative, the work done by this force is equal to the difference of the average values of $\phi_n$,

$$\mathfrak{W} = \langle \phi_n \rangle_{\mathrm{f}} - \langle \phi_n \rangle_{\mathrm{i}} = N \langle \ln F_n \rangle_{\mathrm{f}} - N \langle \ln F_n \rangle_{\mathrm{i}} + \mu N (\langle \Omega_n \rangle_{\mathrm{i}} - \langle \Omega_n \rangle_{\mathrm{f}}) . \tag{S66}$$

Since the final and initial cumulative growth of the population are roughly the same, as shown numerically in Fig. 4(b), we conclude that $\mathfrak{W} \simeq \mu N (\langle \Omega_n \rangle_{\mathrm{i}} - \langle \Omega_n \rangle_{\mathrm{f}}) \simeq 0.1 \times 70 \times 0.2 = 1.4 > 0$. This results shows how the force $\phi_n - \phi_{n'}$ actively contributes to *reduce* the population sensitivity to variations.

#### VI. MATHEMATICAL DETAILS OF *TWO-TYPES MODEL* DISCUSSED IN BOX 2

Here, we elucidate the mathematical expressions behind the two-types model discussed in Box 2.

The *reproduction rate force* of this model can be expressed as

$$\sum_\gamma (n_\gamma - n'_\gamma) \ln f_\gamma = (\text{\# individuals varying from } \gamma_2 \text{ to } \gamma_1) \times \ln \frac{f_{\gamma_1}}{f_{\gamma_2}} \,. \tag{S67}$$

which clarify the intuitively obvious fact that types $\gamma_1$ are favored.

The *entropic force* is nonvanishing only when the composition changes its "degree of homogeneity", *i.e.* $\{(3,0),(0,3)\} \to \{(2,1),(1,2)\}$ (or its reverse), and it is equal to

$$\sum_\gamma \left(\ln n'_\gamma! - \ln n_\gamma!\right) = \ln 3 \,. \tag{S68}$$

The effect of the *nonconservative force* is more idiosyncratic. For $\pi_{\gamma_1 \leftarrow \gamma_2} = \pi_{\gamma_2 \leftarrow \gamma_1} = \mu$, such force can be expressed as

$$\zeta_{(0,3)\leftarrow(3,0)} = 0 \tag{S69a}$$

$$\zeta_{(0,3)\leftarrow(1,2)} = \zeta_{(3,0)\leftarrow(2,1)} = \ln \frac{(2-\mu)^3}{27\mu(1-\mu)^2} \tag{S69b}$$

$$\zeta_{(2,1)\leftarrow(1,2)} = 0 \tag{S69c}$$

$$\zeta_{(3,0)\leftarrow(1,2)} = \zeta_{(0,3)\leftarrow(2,1)} = \ln \frac{(1+\mu)^3}{27\mu^2(1-\mu)} \,. \tag{S69d}$$

For $\mu < 1/2$, this force favors the homogeneous states $\{(3,0),(0,3)\}$. When summed along the cycles depicted in Box 2, this force becomes

$$\zeta_{\text{cycle}} = \zeta_{(0,3)\leftarrow(3,0)} + \zeta_{(1,2)\leftarrow(0,3)} + \zeta_{(3,0)\leftarrow(1,2)} = \zeta_{(3,0)\leftarrow(0,3)} + \zeta_{(2,1)\leftarrow(3,0)} + \zeta_{(0,3)\leftarrow(2,1)} = \ln \frac{(1-\mu)(1+\mu)^3}{(2-\mu)^3\mu} \,. \tag{S70}$$

The force arising from the *variants growth potential* (S10), $\phi_n - \phi_{n'}$ simply favors the population with the highest expected growth at the next generation. For each population composition, we have

$$\phi_{(3,0)} = 3 \ln \left\{ \left[ f_{\gamma_1}(1-\mu) + f_{\gamma_2}\mu \right] 3 \right\} \tag{S71a}$$

$$\phi_{(2,1)} = 3 \ln \left\{ \left[ f_{\gamma_1}(1-\mu) + f_{\gamma_2}\mu \right] 2 + \left[ f_{\gamma_1}\mu + f_{\gamma_2}(1-\mu) \right] \right\} \tag{S71b}$$

$$\phi_{(1,2)} = 3 \ln \left\{ \left[ f_{\gamma_1}(1-\mu) + f_{\gamma_2}\mu \right] + \left[ f_{\gamma_1}\mu + f_{\gamma_2}(1-\mu) \right] 2 \right\} \tag{S71c}$$

$$\phi_{(0,3)} = 3 \ln \left\{ \left[ f_{\gamma_1}\mu + f_{\gamma_2}(1-\mu) \right] 3 \right\} \tag{S71d}$$

For low probability of variations, $\mu < 1/2$, this force favors the population with the highest number of fast-growing types, like the reproduction rate force. Due to the simplicity of the model, the aspect of this force related to robustness cannot be highlighted.

#### VII. CONNECTION WITH THE OTHER MODELS

Our model (S6) can be regarded as a generalization of Wright–Fisher model, which features the multinomial term to describe selection fluctuations, see *e.g.* [7]. In contrast to the latter model, however, we specify neither the nature of $\gamma$—*i.e.* genetic variables such as alleles—, nor the kind of variations—*i.e.* genetic variations like mutations of alleles.

A similar discussion holds for the quasispecies model, which we introduce next.

##### A. Quasispecies Model

The quasispecies model was first formulated by M. Eigen to describe a system of information-bearing self-replicating macromolecules [8, 9]. As it conceptualizes well the dynamics of viral infections, this model is now well established in the context of evolutionary virology [10]. We here review its discrete-time formulation [2, 11], and relate it to our reproduction–variation–selection dynamics.

Quasispecies dynamics describes the evolution of *type frequencies* $z_\gamma := n_\gamma / N$. Since the population size $N$ is taken as infinite, $z_\gamma$ is treated as a continuous variable that evolves deterministically. Over one generation, $\tau \to \tau + 1$, first selection and then variations affect $z_\gamma(\tau)$,

$$z_\gamma(\tau+1) = \frac{\sum_{\gamma'} \pi_{\gamma\leftarrow\gamma'} f_{\gamma'} z_{\gamma'}(\tau)}{\sum_{\gamma'} f_{\gamma'} z_{\gamma'}(\tau)} \,. \tag{S72}$$

Indeed, the factor $f_{\gamma'} z_{\gamma'}(\tau)$ quantifies the reproduction of each type and thus determines the rate at which each type is selected. The term $\pi_{\gamma\leftarrow\gamma'}$ determines the rate of variation $\gamma \leftarrow \gamma'$ over one generation. Finally, the denominator $\sum_{\gamma'} f_{\gamma'} z_{\gamma'}(\tau)$ is the overall growth of the population and ensures the normalization of the frequencies $z_\gamma$.

To show the connection between the quasispecies dynamics (S72) and our stochastic model (S6), we now establish the deterministic limit of the latter. In the limit of large $N$, by applying the Sanov theorem to the kernel (S6) (for a rigorous application see *e.g.* Ref. [12]), we obtain

$$W_{n\leftarrow n'} \simeq \exp\left\{-N\mathcal{D}\left(z_\gamma \middle\| \frac{f_\gamma \sum_{\gamma'} \pi_{\gamma\leftarrow\gamma'} n'_{\gamma'}}{\sum_{\gamma'\gamma''} f_{\gamma''} \pi_{\gamma''\leftarrow\gamma'} n'_{\gamma'}}\right)\right\}, \tag{S73}$$

where

$$\mathcal{D}\left(z_\gamma \middle\| \frac{f_\gamma \sum_{\gamma'} \pi_{\gamma\leftarrow\gamma'} n'_{\gamma'}}{\sum_{\gamma'\gamma''} f_{\gamma''} \pi_{\gamma''\leftarrow\gamma'} n'_{\gamma'}}\right) = \sum_\gamma z_\gamma \ln \frac{z_\gamma}{\frac{f_\gamma \sum_{\gamma'} \pi_{\gamma\leftarrow\gamma'} n'_{\gamma'}}{\sum_{\gamma'\gamma''} f_{\gamma''} \pi_{\gamma''\leftarrow\gamma'} n'_{\gamma'}}} \geq 0 \tag{S74}$$

denotes a Kullback–Leibler divergence. In this limit, $W_{n\leftarrow n'}$ becomes very peaked around the frequency $z = \left(\frac{f_\gamma \sum_{\gamma'} \pi_{\gamma\leftarrow\gamma'} n'_{\gamma'}}{\sum_{\gamma'\gamma''} f_{\gamma''} \pi_{\gamma''\leftarrow\gamma'} n'_{\gamma'}}\right)_\gamma$, and the stochastic dynamics described by the master equation $p_n(\tau+1) = \sum_{n'} W_{n\leftarrow n'}\, p_{n'}(\tau)$ evolves deterministically. Indeed, any population frequency at generation $\tau$, $z_\gamma(\tau)$, is mapped into

$$z_\gamma(\tau+1) = \frac{f_\gamma \sum_{\gamma'} \pi_{\gamma\leftarrow\gamma'} z_{\gamma'}(\tau)}{\sum_{\gamma'\gamma''} f_{\gamma''} \pi_{\gamma''\leftarrow\gamma'} z_{\gamma'}(\tau)} \tag{S75}$$

at the next generation.

In contrast to the quasispecies model (S72), in (S75) the step of reproduction follows that of variations. In addition, quasispecies models differ from RVS dynamics as they are typically restricted to genetic types with sequential genomes of finite length, and they limit variations to point mutations.

#### VIII. DISCUSSION OF EXPERIMENTAL RESULTS: MONTVILLE *ET AL.* EXPERIMENT ON VIRAL EVOLUTION

The two major implications of our fluctuation relation are as follows. *(i)* As the variation rate increases, the sensitivity to variations decreases. *(ii)* As the sensitivity to variations decreases with $\mu$ and approaches a plateau, the fluctuations of sensitivity decrease as well. We here discuss these implications in the context of recent experiments involving viral evolution.

Viruses, and in particular RNA viruses, are ideal to test prediction of simple evolutionary models: their growth is fast and can be quantified; their mutation rate is typically high, $10^{-5}$–$10^{-3}$ variations per nucleotide copied; and their population size can be controlled [10].

In Ref. [6], two sets of three viral populations founded from the same lineage are evolved under strong selection but different effective mutation rates, *low* vs *high*. This difference was achieved thanks to the phenomenon of complementation: when multiple viral particles infect the same host, deleterious mutations affecting one particle may be compensated by the proper functioning of the co-infecting particles. As a consequence, growth is negatively affected by mutations only when multiple viruses are mutated. The probability that this happens is given by the product of the mutation probability of the single viruses. Therefore, by controlling the level of co-infection, one can control the effective probability of mutations $\mu$ that each population experiences: higher the co-infection, lower the effective mutation rate.

To test sensitivity to mutations of the viruses of each population, the authors isolated 10 clones from each of them (hence 30 clones for each of the two levels of co-infection). Mutations accumulated in these clones as they were let evolve under low selection and low population number, a procedure known as *bottlenecking*. In this way, an average of 1.3 mutations accumulated in each virus.

Sensitivity to such mutations can be assessed by comparing the growth of such mutagenized viruses against that of their ancestors, $\Delta \ln F := \ln F_{\mathrm{anc}} - \ln F_{\mathrm{mut}}$. Note that, for low values of $\Delta \ln F_n$, one has $\Delta \ln F_n \simeq \Omega_n$, as shown in Eq. (S65).

We re-evaluated the data reported in Fig. 1 of Ref. [6] (the outlier data point identified by Montville *et al.* is left out of our analysis) and we found that

$$\begin{aligned} \langle \Omega_n \rangle_{\mathrm{l}} &= 0.03 \pm 0.02\,, \quad \mathrm{var}\{\Omega_n\}_{\mathrm{l}} = (1.3 \pm 0.4)\,10^{-2} \\ \langle \Omega_n \rangle_{\mathrm{h}} &= 0.11 \pm 0.03\,, \quad \mathrm{var}\{\Omega_n\}_{\mathrm{h}} = (2.3 \pm 0.6)\,10^{-2}\,, \end{aligned} \tag{S76}$$

which are compatible with our predictions:

$$\langle \Omega_n \rangle_{\mathrm{l}} < \langle \Omega_n \rangle_{\mathrm{h}} \tag{S77}$$

$$\mathrm{var}\{\Omega_n\}_{\mathrm{l}} < \mathrm{var}\{\Omega_n\}_{\mathrm{h}} \,. \tag{S78}$$

However, as mentioned in the main text, these results are not fully statistically significant. When testing the hypothesis (S77), we found $p = 0.011$ ($t$-test), which is not significantly smaller than the typical 0.05 threshold. Regarding (S78), we found $p = 0.08 > 0.05$ (Fisher Ratio test), which does not allows us to reject the null hypothesis. We thus conclude that, although the results are promising, further experimental data are needed to test the inequalities (S77) and (S78).

[1] J. H. Gillespie, *Population genetics: a concise guide* (The Johns Hopkins University Press, Baltimore (MD), 2004).
[2] L. Peliti, arXiv **cond-mat/9712027v1** (1997), cond-mat/9712027v1.
[3] D. Simon, *Evolutionary Optimization Algorithms* (John Wiley & Sons, 2013).
[4] P. Schuster and J. Swetina, Bull. Math. Biol. **50**, 635 (1988).
[5] E. van Nimwegen, J. P. Crutchfield, and M. Huynen, Proc. Natl. Acad. Sci. U.S.A. **96**, 9716 (1999).
[6] R. Montville, R. Froissart, S. K. Remold, O. Tenaillon, and P. E. Turner, PLoS Biol. **3**, e381 (2005).
[7] W. J. Ewens, *Mathematical Population Genetics I. Theoretical Introduction* (Springer, New York (NY), 2004).
[8] M. Eigen, Naturwissenschaften **58**, 465 (1971).
[9] M. Eigen and P. Schuster, *The Hypercycle* (Springer, 1979).
[10] E. Domingo, J. Sheldon, and C. Perales, Microbiol. Mol. Biol. Rev. **76**, 159 (2012).
[11] D. B. Saakian and C.-K. Hu, in *Current Topics in Microbiology and Immunology* (Springer, 2015) pp. 121–139.
[12] H. Touchette, Phys. Rep. **478**, 1 (2009).
